# Wide and Deep Imaging of Neuronal Activities by a Wearable NeuroImager Reveals Premotor Activity in the Whole Motor Cortex

**DOI:** 10.1101/434035

**Authors:** Takuma Kobayashi, Tanvir Islam, Masaaki Sato, Masamichi Ohkura, Junichi Nakai, Yasunori Hayashi, Hitoshi Okamoto

## Abstract

Wearable technologies for functional whole brain imaging in freely moving animals would advance our understanding of cognitive processing and adaptive behavior. Fluorescence imaging can visualize the activity of individual neurons in real time, but conventional microscopes have limited sample coverage in both the width and depth of view. Here we developed a novel head-mounted laser camera (HLC) with macro and deep-focus lenses that enable fluorescence imaging at cellular resolution for comprehensive imaging in mice expressing a layer- and cell type-specific calcium probe. We visualized orientation selectivity in individual excitatory neurons across the whole visual cortex of one hemisphere, and cell assembly expressing the premotor activity that precedes voluntary movement across the motor cortex of both hemispheres. Including options for multiplex and wireless interfaces, our wearable, wide- and deep-imaging HLC technology could enable simple and economical mapping of neuronal populations underlying cognition and behavior.

## 1. Introduction

Wearable imaging instruments represent an emerging class of powerful and versatile measurement tools for *in vivo* functional analysis of the brain in freely moving animals^1–4^. Wearable microscopes such as head-mounted 2-photon microscopes, miniature endoscopes, fiber photometers, and other implantable devices have already made significant contributions in neuroscience^5–9^. Many of these instruments also incorporate recent improvements in fluorescent probe technology, such as the genetically encoded Ca^2+^ indicators^10–13^, which enable long-period, real-time imaging of neuronal activity at high signal-to-noise ratios in the living animal brain. Despite such advances, wide-field imaging of cortico-cortical inter-regional interactions at high spatiotemporal resolution remains difficult. Most current instruments only allow the capture of images from a single or limited number of focal planes. Also, because ongoing animal movement often breaks the instrument, continuation of the experiment can be difficult if a device is too delicate and expensive to obtain replacements. To address these problems, we developed a novel wearable fluorescence imaging system containing a head-mounted laser camera (HLC) with deep-focus and macro photographic lenses that can comprehensively analyze neuronal activity in the mouse cerebral cortex.

Traditionally, researchers have attempted to improve microscopic optics to obtain wider and deeper fields of view under the restricting design conditions of a defined focal plane and optical aberration correction. Here, we tried to achieve the same goal, but with an alternative approach using a deep-focus optical system that can integrate images of objects at different depths and perspectives into a single-plane image. This apparatus adopts an optical system similar to the one used in inexpensive compact cameras and smart cellular phones, thereby allowing us to obtain images similar to the z-stack images of multiple focal planes produced by two-photon laser microscopy. Many types of camera modules are commercially available, thus we could manufacture a compact HLC imaging system on a purpose-built or mass-production commercial scale, and within a reasonable budget.

In this study, we used a hand-made HLC for *in vitro* and *in vivo* fluorescence Ca^2+^ imaging to assess whether physiological neuronal activity could be visualized over a wide view in the deep cortical layers of a freely moving mouse. We also show that we can resolve the information of activities of individual cells by application of proper image processing algorithm. First, we imaged individual neurons with orientation selectivity in the visual cortex to analyze cellular physiology, and second, we applied the HLC to identify the cellular assembly used for premotor activity during the planning phase before the initiation of voluntary movement in the motor cortex of both hemispheres.

## 2. Results

### 2.1 Development of a wearable instrument for fluorescence imaging of neural activity in the cerebral cortex

Figure 1 demonstrates our wearable fluorescence imaging system developed to visualize neural activity in the cerebral cortex of freely moving mice (left panel in Fig. 1a, b, Methods section; surgical method is explained in Supplementary Fig. 1a). The wearable apparatus consists of a separable camera and a spacer with a cranial window at its base. Imaging of the cortex is performed by the camera component through the cranial window. The object plate of the cranial window makes direct contact with the surface of the cortex (Fig. 1b). Repeated imaging of a specific brain area can be easily carried out while housing the mouse for a long period (Supplementary Fig. 1b). By equipping a suitable excitation light source and an absorption filter, the HLC can perform imaging for green or red fluorescence (right panels in Fig. 1a). The HLC is compact and lightweight, and therefore does not impede an animal’s normal behavior (Fig. 1c, Supplementary Movie 1). We confirmed that the imaging of neuronal activities by the HLC attached to the head of freely moving mice causes no significant increase of stress, and does not affect the locomotor activity and the behavioral pattern (Supplementary Fig. 2).

**Figure 1.**
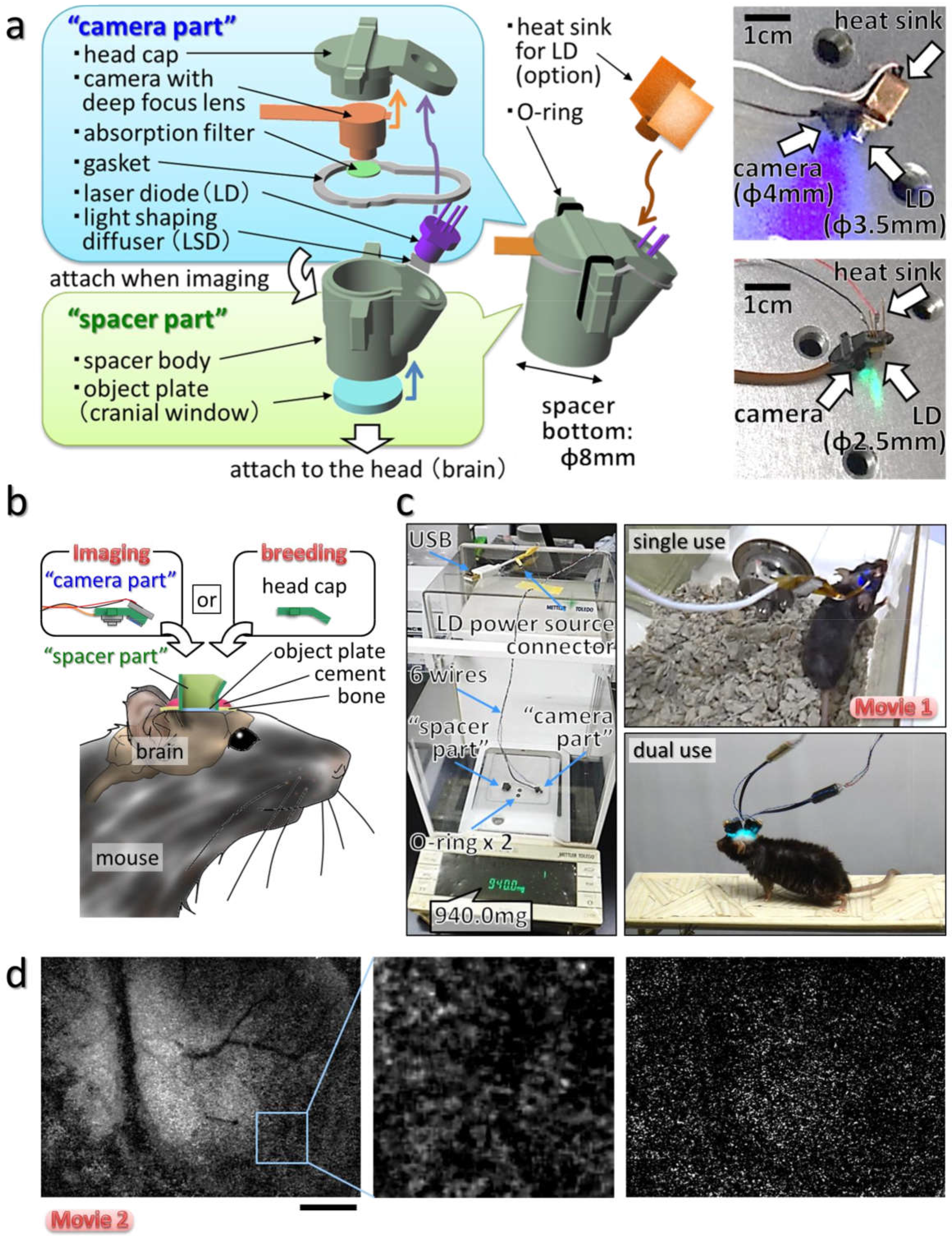
Wearable system for imaging the cerebral cortex in freely moving animals **(a)** Schematic image of the head-mounted laser camera (HLC) is shown. Right photos show the HLCs for green (GFP) or red fluorescence (RFP) imaging. **(b)** Schematic image of the HLC imaging system. The camera part is attached to the spacer part when imaging is performed, and the head cap is attached to the spacer part when the mouse is housing. **(c)** The weight measurement of all the constituent parts of a typical HLC is shown in the left image. Freely moving mice equipped with a single or dual HLC’s are shown in the right column images (Supplementary Movie 1). **(d)** Fluorescence Ca^2+^ imaging of the occipital cortical area including the visual cortex on an awake CaMK2a-G-CaMP mouse using the HLC (see also Supplementary Movie 2). The left image is a representative single frame of the movie. The middle image is a magnified view of the inset in the left image. The right image is a representative frame of the subtracted imaging movie, which was made by subtracting the average of the FI of each pixel from the FI of the same pixel during the first 10 frames of the imaging movie. The HLC can visualize excitatory neuronal activity as blinking light spots, and also visualize the spatial distribution of fine capillaries with a wide field of view (4.25 × 5.66 mm). The view area of the HLC is adjustable. Bar indicates 1 mm.

Additionally, multiple cameras can be attached to the heads of small mice such as the C57BL/6 line (Fig. 1c, Supplementary Fig. 1c-e) to enable the simultaneous imaging of multiple loci, which previously has proved difficult. Importantly, since the wearable HLC is robustly fixed to the mouse skull, normal body movements do not perturb the wide-view fluorescence imaging (Fig. 1d, Supplementary Movie 2). The number of blinking spots representing presumptive cellular elements in the field of view in Fig. 1d was estimated at approximately 10,000 (Supplementary Fig. 3, Supplementary Movie 3).

We used a laser diode (LD) in the HLC as an excitation light source. Thus, to prevent possible temperature increases in the diode due to continuous lighting during long-term imaging, we tested an LD driver with an attached pulse generator to toggle the LD on and off (Supplementary Fig. 4a) and heat sinks of different sizes. This modified design seemed to effectively suppress temperature changes in the LD that could adversely affect the neuronal imaging (Supplementary Fig. 4b-f). Furthermore, pulse driving strengthens the laser light, and such an integrated light source makes the system more adaptable to wireless control than one dependent on an external light source via optical fiber (Supplementary Fig. 5). Therefore, the HLC imaging system is suitable for use on constantly moving animals by using either wired or wireless transmission because the number of output channels is small enough to allow a USB connection and low electric power consumption with a CMOS image sensor.

Before moving to *in vivo* imaging, the *in vitro* optical specifications of the HLC were investigated (Fig. 2a-f, Methods section). A checkerboard chart (Fig. 2a, b) or a line chart (Fig. 2c-f) was captured by the HLC, and each optical parameter was calculated based on the actual measurement values. As a result, a maximum spatial resolution 4.17 μm/pixel was obtained if the blurring of edges in the images were ignored (Fig. 2a). The optical distortion was 2.78% at maximum (television distortion = 3.91%), and the depth of field exceeded 19.5 mm (see Methods section for details of the calculations).

**Figure 2.**
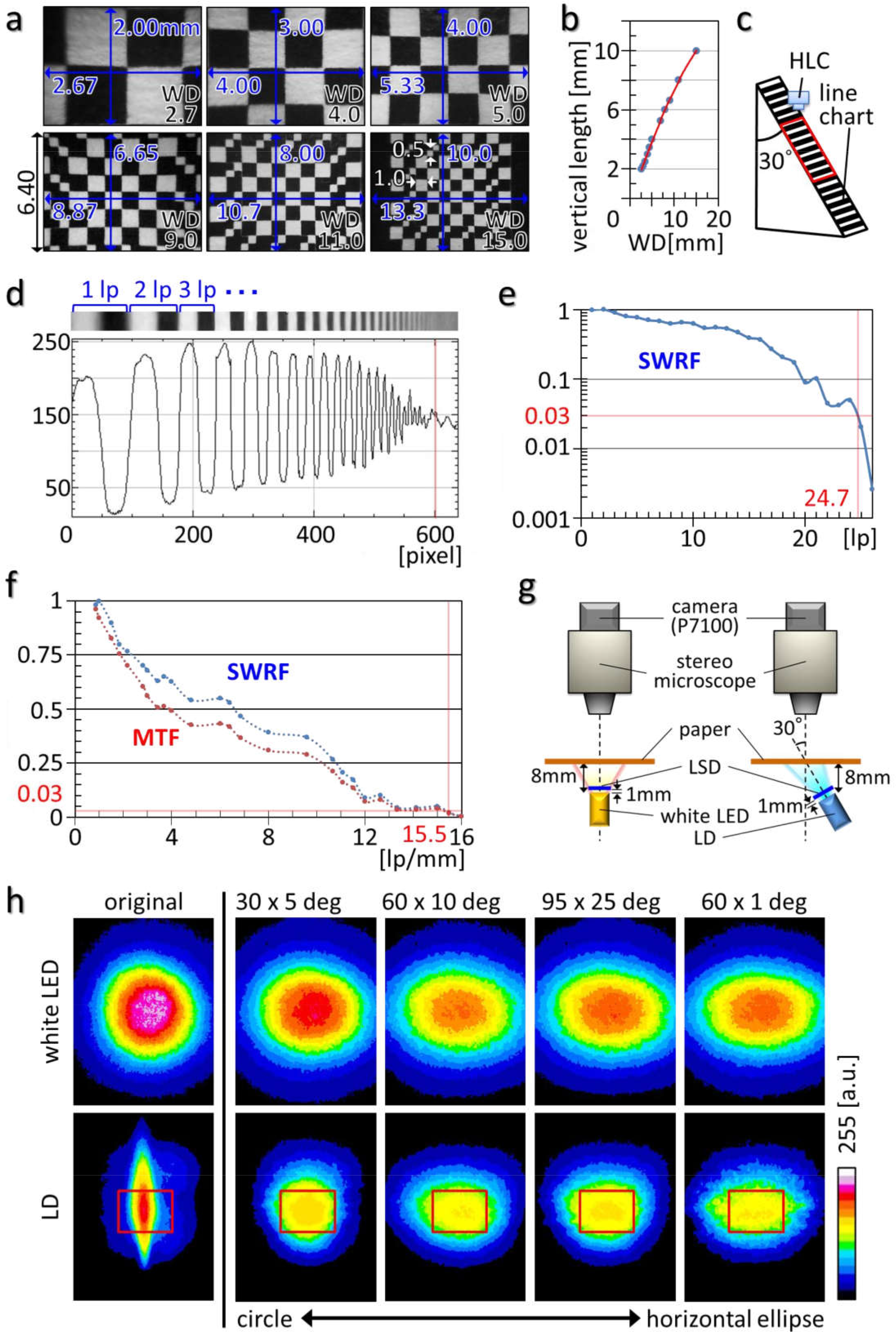
Optical specifications and adjustments of the HLC **(a, b)** A checkerboard-design chart was captured by the HLC using different fields of view. All results are shown in (b), using the representative images shown in order of size of view from (a). The red line in (b) indicates an approximate curve, expressed by Y = −1.62 × 10^−2^X^2^ + 9.4X − 0.48, and the correlation coefficient is R^2^ = 0.999. **(c)** Apparatus used for measuring a line-spread function (LSF) of the HLC using a line chart. The red square indicates the view area of the HLC. **(d)** The upper picture shows the image of the line chart taken by the HLC, and the lower graph indicates the LSF (ordinate, light intensity; abscissa, vertical position of the image). The red line corresponds to the position of the red lines in (e, f). **(e, f)** A square wave response function (SWRF) was calculated based on the LSF (e) (lp = line pair, for a detailed calculation process see Methods section). A modulation transfer function (MTF) was calculated based on the SWRF (f). The horizontal and vertical red lines were drawn to cross at the point where the SWRF = 0.03. **(g)** Schematic image representing the light shape measurements. Several types of light-shaping diffusers (LSDs) were attached in front of the white LED or LD at a distance of 1 mm. **(h)** The distributions of illumination by an LD, whose original beam divergence was 3.3 × 14 deg at 40 mA, are shown as an 8-bit pseudo-colored image. The distributions of illumination by a white LED are shown as controls. Numbers at the top of the images show the LSD characteristics. Images are arranged according to the order of deviation of the LSD from the circle to the horizontal ellipse. Red squares indicate the typical view area of the HLC (4.0 × 5.3 mm).

Next, the optical processing conditions for homogenizing the flat irradiation shape of a coherent laser beam emitted by the LD were examined using a light-shaping diffuser (LSD) (Fig. 2g, h, Supplementary Fig. 4g, h). Based on actual measurements using various types of LSD (Fig. 2h) and comparison with the theoretical value with simulation (Supplementary Fig. 4g, h), a 60 × 10 degree LSD was found to be the most preferable. Accordingly, we expected that the HLC equipped with a deep-focus optical system could capture images from wide brain areas and various depths at quasi-cellular resolution.

To verify the imaging ability of the HLC, *in vitro* and *in vivo* fluorescence-imaging tests were performed. First, images of the fluorescent beads implanted into the cerebral cortex of the mouse were compared between the HLC and a conventional stereomicroscope (Fig. 3a-c). The HLC, but not the conventional stereomicroscope could detect beads of similar size to cells in the deep cortex even at 800 μm in depth (Fig. 3d, e). Additionally, the deep-position HLC images showed less blurring of the bead shapes than those taken by stereomicroscope (Fig. 3f). The findings indicate that the HLC can acquire fluorescence signals deep in the cerebral cortex at high precision compared to a conventional wide-field fluorescence microscope. Second, the Ca^2+^ imaging was performed with 3D cultured cells made by embedding transfected Hela cells in an extracellular matrix gel as a mock brain tissue (Fig. 4a). The change ratios (ΔF/F) of the fluorescence intensity (FI) of individual cells at various depths increased with histamine administration, whereas they were decreased by EGTA administration (Fig. 4b, Supplementary Fig. 6a, Supplementary Movie 4). The results indicate that the Ca^2+^ imaging of individual cells could be performed in a 3D cell culture using the HLC.

**Figure 3.**
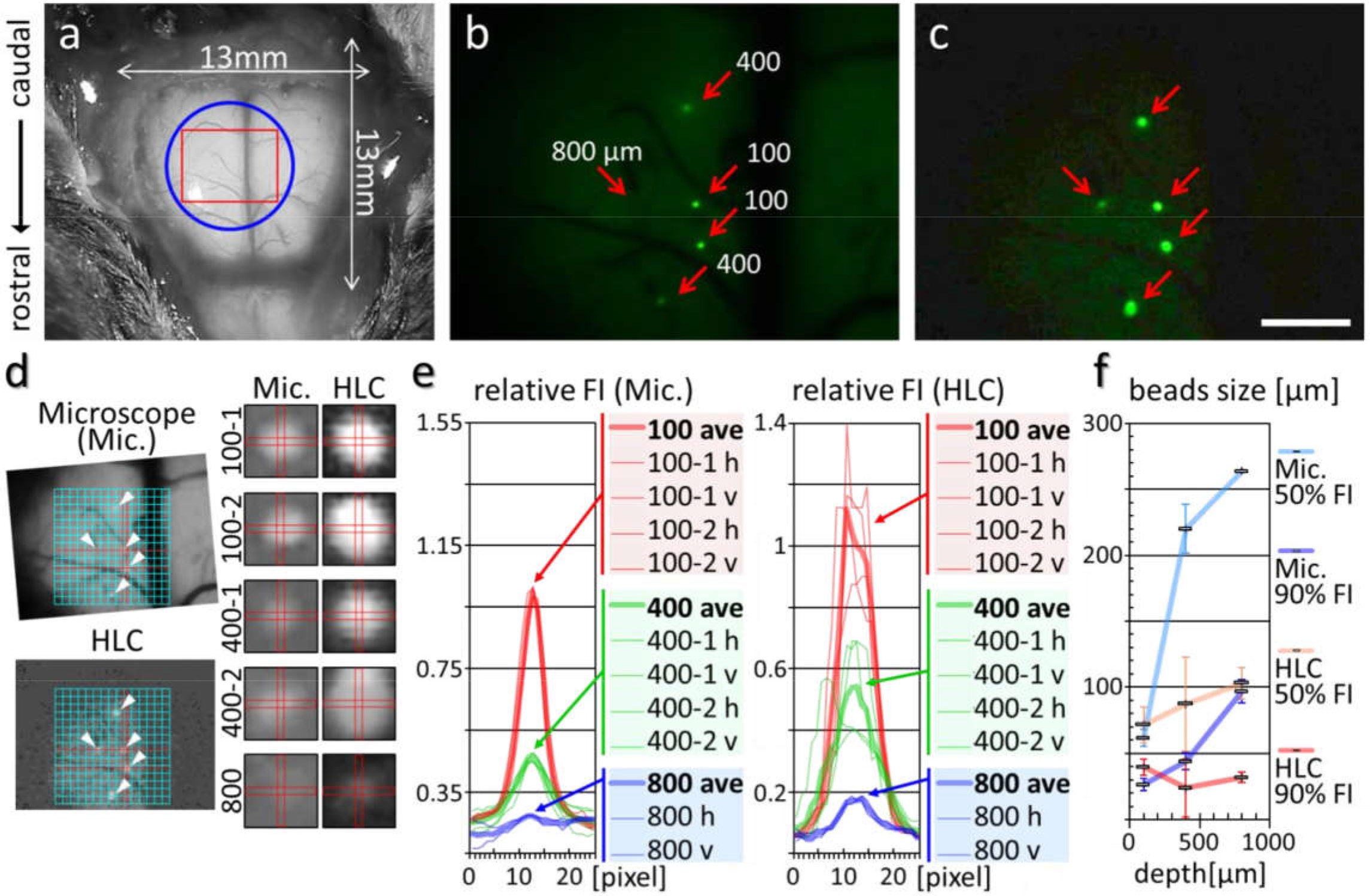
The HLC can acquire the fluorescence signal of the beads implanted into the deep layers of the cerebral cortex **(a)** Fluorescent beads were implanted into the mouse cortex after a craniotomy. The blue circle and red square indicate the spacer position and the imaging area of the HLC, respectively. **(b, c)** Fluorescence images taken by a fluorescence stereo microscope (b) and the HLC (c). Red arrows indicate the positions of implanted 15-μm beads, and the numbers indicate the depth of implantation. Bar indicates 1 mm. **(d)** In the two left images, the size and position of (b) and (c) were aligned based on the pattern of blood vessels and the central position of the beads implanted at a depth of 100 μm (red lines). White arrowheads indicate the bead positions. The right two rows of 10 images show the magnified images of each bead taken by the stereo microscope (Mic., left column) or the HLC (right column). The vertical (v) and horizontal (h) rectangles were drawn to cross at the center of each bead. **(e)** Fluorescence intensity (FI) distribution along the (v) and (h) in (d) were measured, and the relative FIs against averaged FI at the center of the bead at a 100-μm depth are shown. **(f)** A summary of (e) is shown. The image size of the fluorescent beads was calculated according to the number of pixels in the image. By comparing the FI at the centers and perimeters of each light spot, the number of pixels of > 50% FI and > 909 FI were counted. In the graph, error bars indicate the standard deviation. Note that the values > 90% of the HLC stay almost invariant regardless of depth (red line).

The spacer part of the device with the cranial window is also useful to observe the cortex under a conventional or 2-photon microscope system (Supplementary Fig. 7, Supplementary Movie 5); however, a major advantage of the HLC in this regard is that a large amount of information can be obtained from a single image in real time compared to scanning multiple focal planes as needed for the same results by confocal or 2-photon laser microscopy systems. Furthermore, by using transgenic animals expressing the Ca^2+^ indicator fluorescent proteins in layer- or cell type-specific manners, we also obtained images from defined subpopulations of neurons in the brain regardless of their distribution areas. As two examples in the present study, CaMK2a-G-CaMP7 mice expressing G-CaMP specifically in excitatory neurons (Fig. 5, 6, Supplementary Fig. 7-9) and Thy1-G-CaMP7 mice expressing G-CaMP predominantly in layer 5/6 pyramidal neurons were imaged (Fig. 4c, d, Supplementary Fig. 6c).

**Figure 4.**
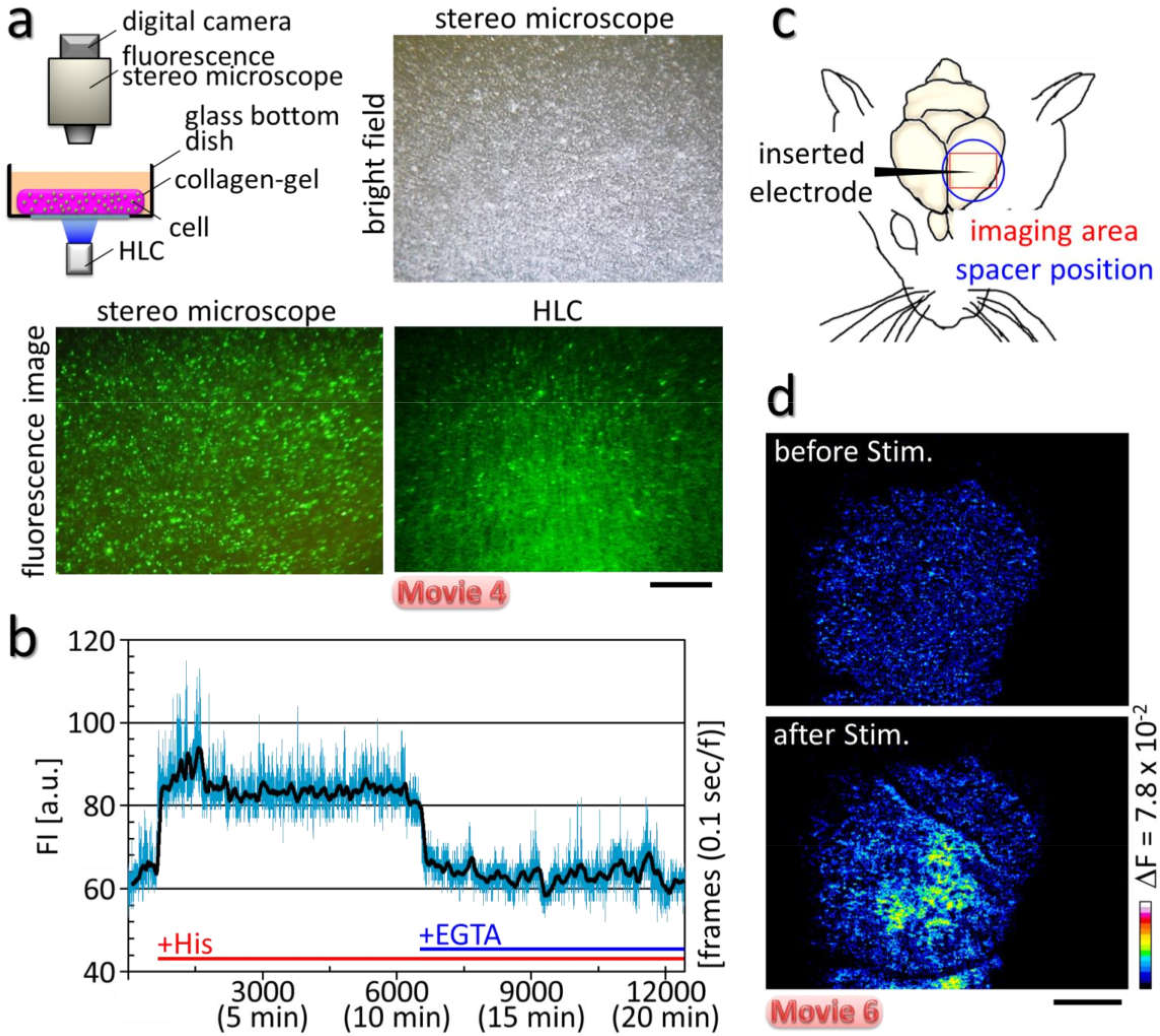
The HLC visualized intracellular Ca^2+^ dynamics of individual cells in a 3D culture, and evoked neuronal activity in the deep layers of cerebral cortex **(a)** Schematic image of the *in vitro* experiment. The sheet of 3D cultured Hela cells transfected with the G-CaMP6 gene was placed into a glass bottom dish, and Ca^2+^ imaging was performed with the HLC. Brightfield and fluorescence images were taken by a stereomicroscope or the HLC. The transfected cells emitted green fluorescence and were distributed sparsely across various depths. Bar indicates 1 mm. **(b)** The graph shows the changes in FI of a single cell. The FI increased after histamine administration (+His), and then decreased when a solution of EGTA was applied. The black line indicates the mean FI for each of 5 frames. **(c)** Schematic diagram for the *in vivo* experiment. An electrode was inserted into the cortex of a Thy1-G-CaMP7 mouse from the side after craniotomy. Then, the HLC was applied to the cortex and Ca^2+^ imaging was performed. **(d)** Pseudo-color images are shown for the change in FI before and after electrical tetanic stimulation that was applied by an inserted electrode (see Supplementary Fig. 6c for more detail). Bar indicates 1 mm.

For Ca^2+^ imaging of the Thy1-G-CaMP7 mouse by the HLC, we applied an electrical tetanus stimulation to the somatosensory area, and observed an increased FI around the electrode in the transgenic mouse, whereas no such evoked signals were detected in a control wild-type mouse (Fig. 4c, d, Supplementary Fig. 6c, Supplementary Movie 6). In Supplementary Movie 6, the HLC was placed on the surface of the cortex without fixing it to the cranial bone. Therefore, the image shows slight vibrations due to the muscle movement induced indirectly by stimulation of the somatosensory area, even under anesthesia. The imaging result indicates that HLC can visualize the evoked activity of cortical neurons located as deep as layer 5/6.

### 2.2 The HLC can detect physiological neuronal activity at cellular resolution in the cerebral cortex including the visual area of one hemisphere in freely moving mice

To demonstrate the validity of the HLC, physiological responses of individual neurons during visual perceptual information processing were observed by Ca^2+^ imaging in freely moving CaMK2a-G-CaMP7 mice (Fig. 5-7).

**Figure 5.**
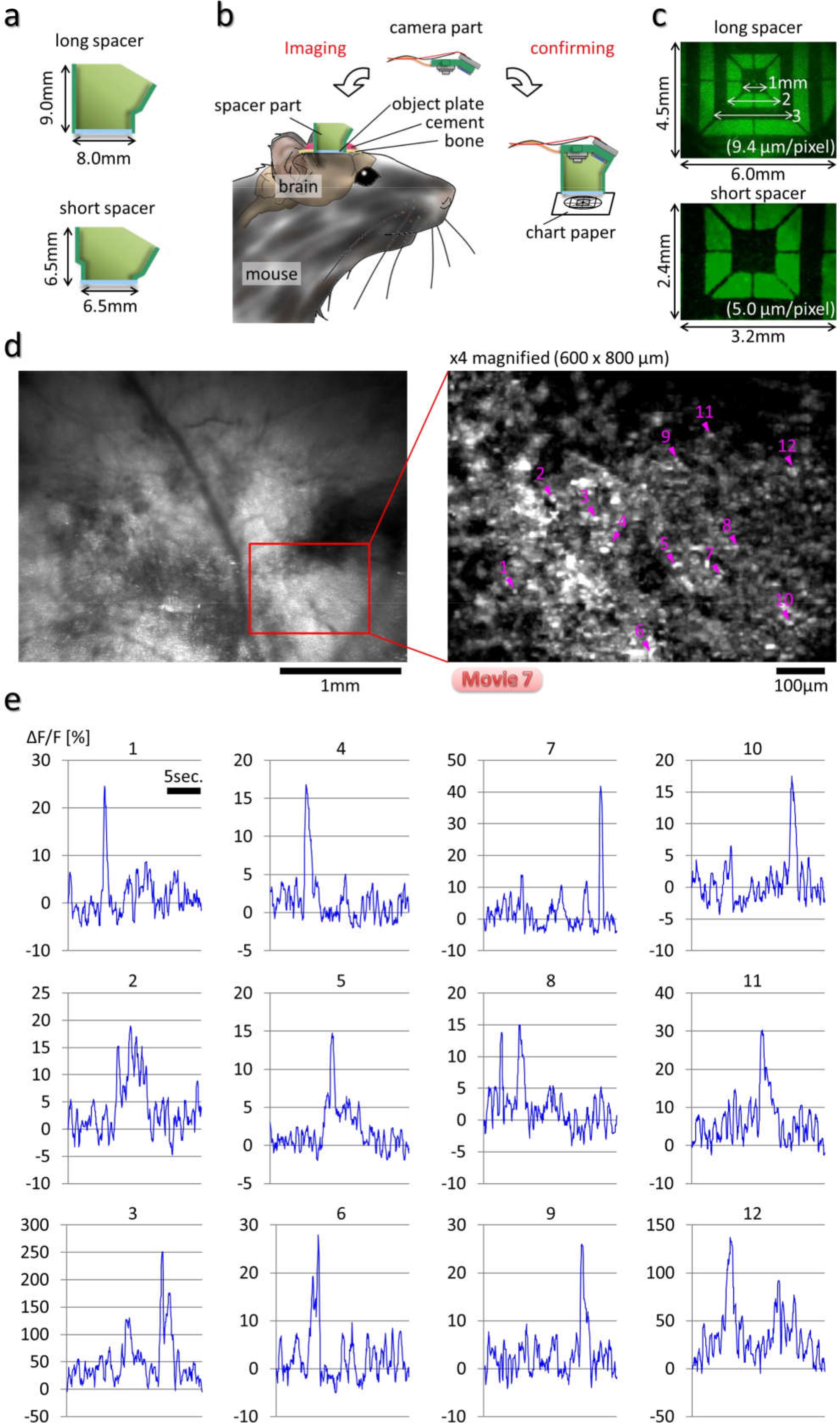
Zoom-up Ca^2+^ Imaging by using the HLC with short spacer in the freely moving mice. The field-of-view size by the HLC can be changed for zooming up by twisting the lens barrel. For small field of view with short working distance after zooming up, it is preferable to use short spacer apparatus. **(a)** Schematic images of different type of spacer are shown. The spacer part can be manufactured arbitrarily by changing its design. **(b)** After imaging, the field-of-view size was confirmed by capturing the image of the reference chart. **(c)** The images of the reference chart captured by the HLC with long or short spacer are shown. **(d)** Ca^2+^ imaging was performed at the visual cortex of the CaMK2a-G-CaMP7 mouse by the HLC with the short spacer under the freely moving condition. The maximized fluorescence image of 20 fps movie for 5 min is shown at the left panel, and its magnified, maximized fluorescence image of the inset part of the left panel for 20 sec is shown on the right panel. Magenta arrowheads and numbers indicate the ROIs of which the light spots were randomly selected. The averaged and maximized movie in each 5 frames is shown in Supplementary Movie 7. **(e)** The change rates of the fluorescence intensity (ΔF/F) at each ROIs of (d) are shown.

By twisting the lens barrel of the camera part of the HLC, the HLC can obtain either a narrow or wide field of view arbitrarily (Fig. 2a, b, Methods section). A short spacer was used to compensate for the reduction in the working distance by optical zooming (Fig. 5a-c). Under the freely moving condition, Ca^2+^ imaging was performed on the visual cortex of the CaMK2a-G-CaMP7 mouse (Fig. 5d, e, Supplementary Movie 7). As a result, the neuronal activity of the visual cortex can be visualized when the mouse was viewing the surrounding scenery, and the vibration derived from mouse’s behavior was not observed and the view field did not drift because of the firm attachment of the HLC to the head. It can be noted that the detection of cells with partial overlap in XY plane but separated in Z axis can raise the possibility of simultaneous recording of cells from various layers of the cortex. To address this problem, we deconvoluted the signals of overlapping cells individually by using a NMF (Non-negative Matrix Factorization) algorithm^14^ (Fig. 6, Methods section). As a result, 685 cells and their activities were detected in the magnified 600 × 800 μm image (Fig. 6c, d). The HLC captures signals from cells at various depths, superimposed in one plane. Though it is not possible to exactly determine the depth of the cells from the surface with the current HLC, we can roughly assume that cells with higher baseline activities may be located on shallower depth from the surface and reconstruct the putative distribution of cells in 3D (Fig. 6e, f).

**Figure 6.**
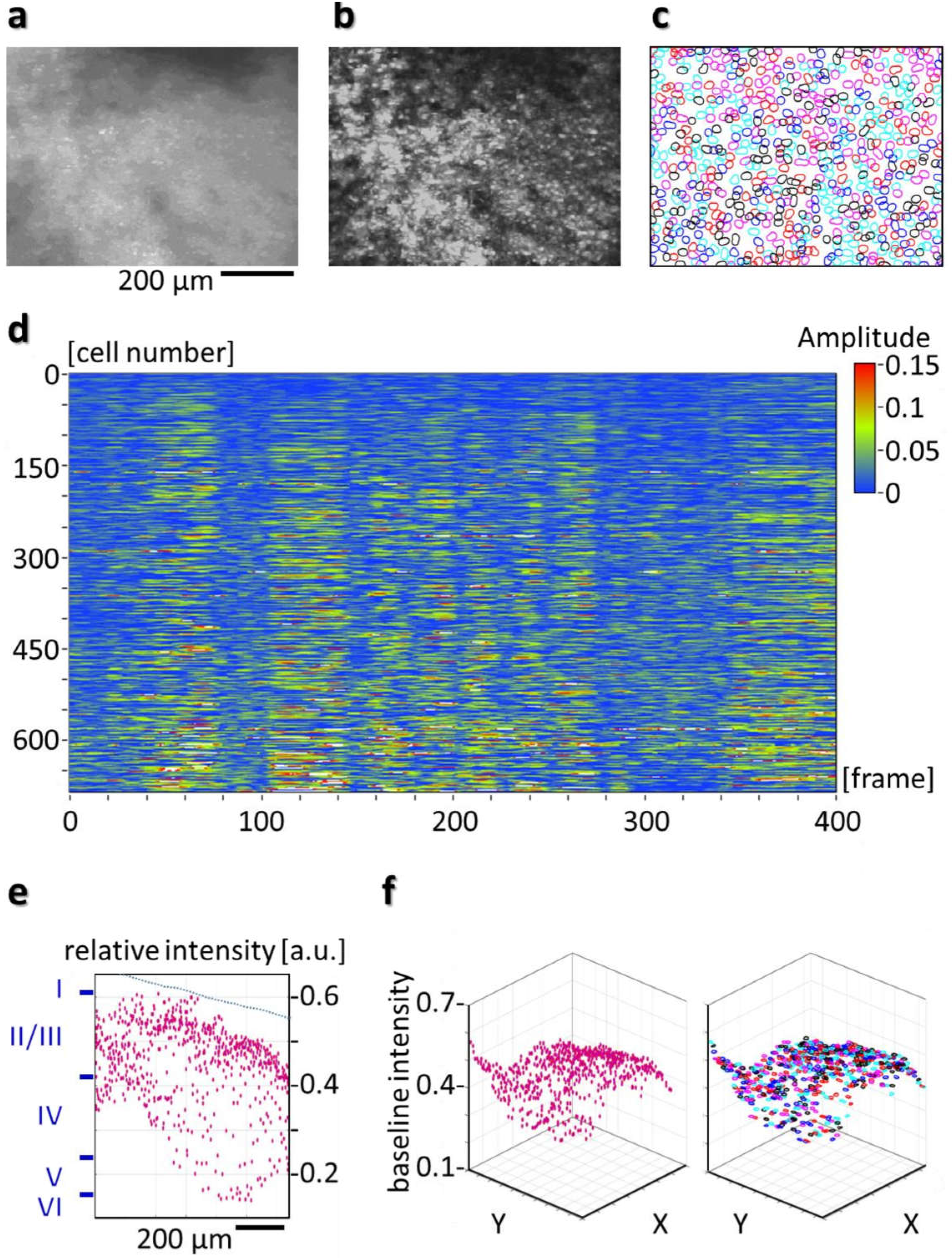
Separation and identification of superimposed signals of individual cells by NMF in the HLC image **(a)** Ca^2+^ imaging of the CaMK2a-G-CaMP7 mouse at the visual cortex was performed under the freely moving condition. The Maximized fluorescence image at 20 fps for 20 sec at the same view point as in Fig. 5d is shown. **(b)** shows the subtracted maximized image of the movie. **(c)** Using the raw 20 fps imaging data for 20 sec (total 400 frames), it was possible to segment the image into individual ROIs (region of interest) representing cell bodies by using a NMF algorithm (Methods section). The result is shown in (c), which shows the boundaries of 685 detected cells, plotted with different colors. It can be noted that many cells with spatial overlap were detected. **(d)** shows the signal extracted by NMF for each cell over time, showing spontaneous activities of cells during free moving. With the application of the NMF algorithm, baseline removal and separation of temporal signals corrupted by spatial overlap of cells was achieved. **(e)** shows the centroid locations of the cell bodies in X-Z plane, where X is the longer image axis of (a) and Z is the depth of cortex corresponding to baseline value of the cell signal. Blue Roman numerals at the Z axis indicate the estimated cortical layer, assuming that deepest signal came from a depth of 1 mm. The broken blue line indicates the imaginary inclined cortical surface. **(f)** The left image shows the centroid locations of the cells in three-dimension, where X and Z axis are same as (e), while Y-axis corresponds to the shorter image axis of (a). A three dimensional distribution of the cell bodies shows the presence of cells with various depths. The right image shows the same three dimensional distributions of the cell bodies with cell boundaries showed instead of centroid.

Next, to examine the differences in physiological responses to different stimuli, a 0.1-second (sec) single flashlight stimulation or 10 repeats of a 0.5-sec flashlight stimulus (light on, 25 ms; light off, 25 ms; 10 times), was applied to the mouse by using an LED positioned in front of the left eye (Methods section). Transient increases (av. 0.50 sec, SD ± 0.14) in FI were observed at 12 fluorescence spots in the primary visual cortex (V1) when a flash stimulation was applied (Fig. 7b), whereas gradual increases in the FI of 7 spots were observed over a longer period (av. 1.09 sec., SD = ± 0.41) during and after repeated flash stimuli (Fig. 7c). The ΔF/F at places other than the visual area, considered to represent basal brain activity irrelevant to visual information processing, showed a fluctuation within 3% maximal FI (black line in the graph). No significant increases of the ΔF/F at these places (black line) during the light stimulus mean that no external light was incident on the imaging area and supports the correctness of the experimental results. Evidently, the transient, significant increases of fluorescence in the visual cortex were caused by neuronal activity, and such increases were strong and long depending on the stimulation time. These results demonstrate that the HLC can differentiate the neuronal responses to different stimuli.

**Figure 7.**
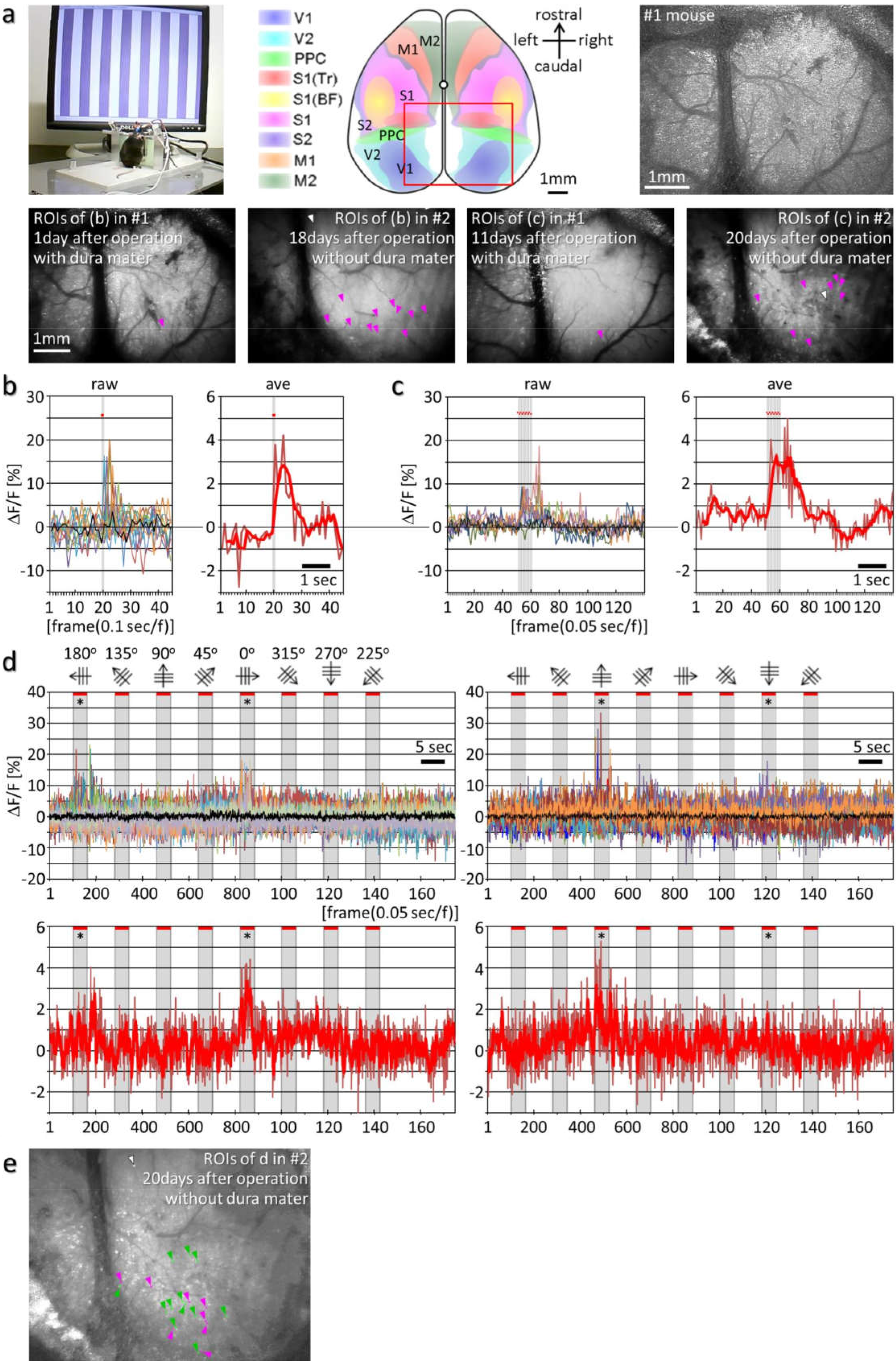
Observation of the physiological responses of individual neurons by large-scale imaging including the whole visual area of the cortex **(a)** The left picture shows the experimental setup. The middle schematic diagram shows the HLC imaging area (red square, 4.25 × 5.66 mm) with the map of the brain area reconstructed from serial sections of the brain atlas (Supplementary Fig. 8a). Abbreviations: V1/2, primary/secondary visual cortex; PPC (PtA), posterior parietal cortex (parietal association cortex); S1 (Tr/BF)/S2, primary (trunk/olfactory barrel field)/secondary somatosensory cortex; M1/2, primary/secondary motor cortex. The right image is the deconvoluted fluorescence image of the occipital cortex in the awake CaMK2a-G-CaMP7 mouse taken by the HLC (see Methods section for detail). The lower four images are fluorescence images of the occipital cortex. Magenta and white arrowheads indicate the region of interests (ROIs) or negative control ROIs, and their ΔF/F are presented in (b), (c). **(b, c)** The left and right graph shows the raw or averaged ΔF/F after a single (b) and 10x repeated flashlight stimulus (c), respectively, was applied by the LED. Gray vertical lines and red dots in each graph indicate the stimulation points. The red bold line data in the right graph of (b) and (c) indicate the means of 3 and 5 frames, respectively. Black lines mean negative control ROIs. **(d)** Drifting gratings in 8 different directions were presented. The upper and lower graphs show the raw or averaged ΔF/F. Examples of the specific ΔF/F increases according to the opposite degree stimulation (asterisks) are shown. Data indicated by the black line indicate the ROI of the negative control, and the gray shaded time windows in each graph indicate the stimulation periods. The red bold line data in the lower graphs indicate the means of 5 frames. **(e)** Fluorescence image shows the ROIs of the #2 mouse. White arrowheads indicate negative control ROIs, and green and magenta arrowheads show ROIs for left graph or right graph in (d). **(f)** shows the distribution of the ROIs of different orientation specificity by different pseudo-colors in the same area as (e). See Methods for details. **(g)** shows the selected ROIs with high level of orientation specific responses (top 3.5 %) among the ROIs in the same area as (e). See Methods for details. These ROIs are mostly contained in the visual area of the cortex. Abbreviations: V1m/b, primary visual cortex monocular/binocular zone; A, anterior; AM, anteromedial; PM, posteromedial; RL, rostrolateral; AL, anterolateral; LM, lateromedial; LI, laterointermediate; POR, postrhinal; P, posterior area.

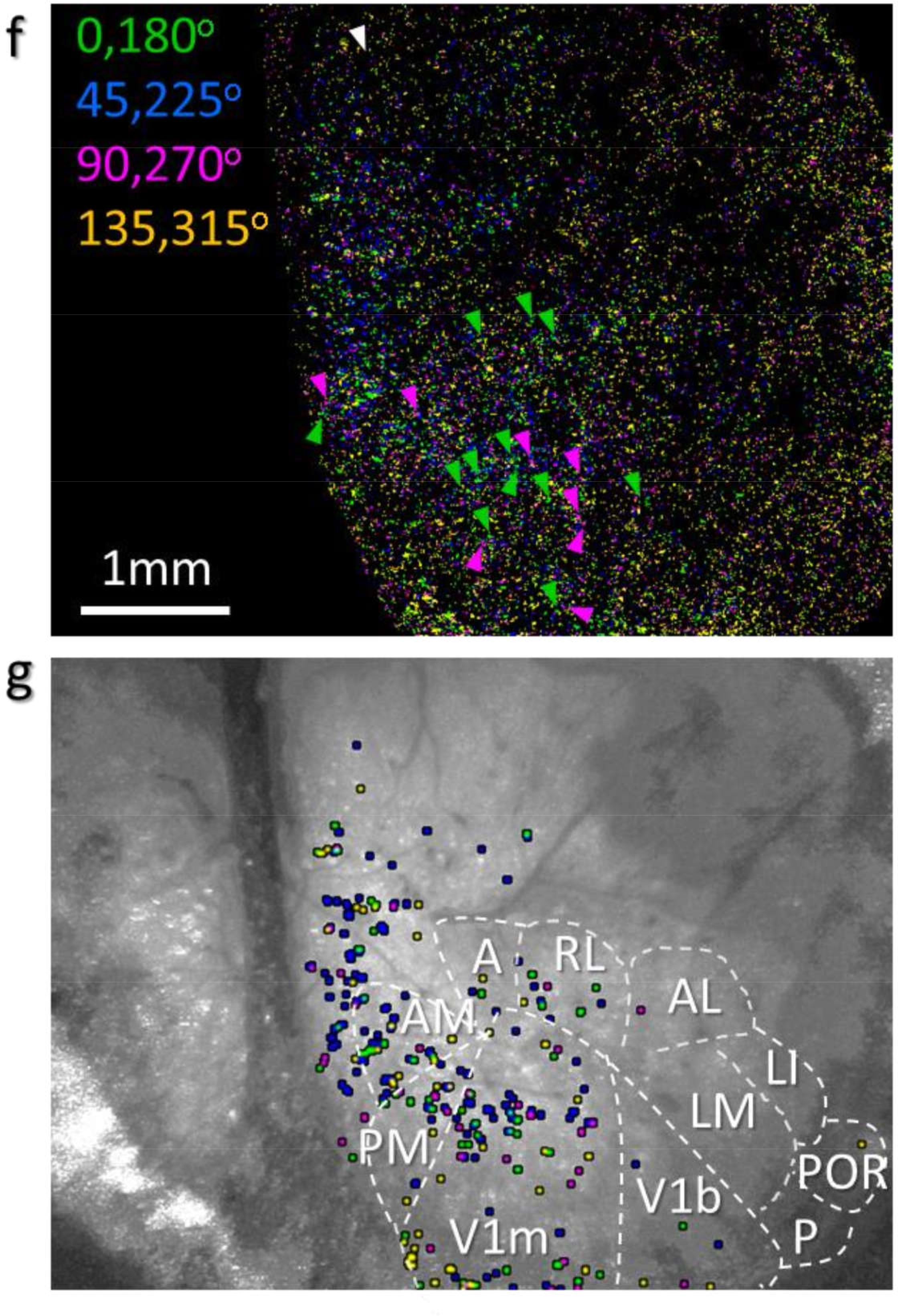

The complex cells in the primary visual cortex in cats show specific orientation selectivity in response to the specific directions of scanning light stimuli^15^. In rodents, the orientation selectivity is represented mainly in layer 2/3 neurons, although minor responses are also observed in the geniculate afferent fibers^16,17^. The neurons with orientation selectivity are also randomly distributed in the visual cortex of rodents, thus we examined whether the HLC could detect such neuronal activity within the visual cortex of mice.

An 8-direction drifting grating was presented to the mice, and Ca^2+^ imaging was performed (Fig. 7a, Methods section). A transient, strong increase in FI was detected at the specific light spots in V1 only when a specifically orientated grating was presented (Fig. 7d). The left column graphs indicate the examples of the 15 light spots that responded robustly to 0 and 180-degree stripes (asterisks), and did not respond at all to 90 and 270 degrees. In contrast the right column graphs indicate the examples of the 12 light spots that responded robustly to 90- and 270-degree stripes (asterisks), while none of them responded to 0 and 180 degrees. We further analyzed the orientation specificity of all ROIs of the observed area (Fig. 7e-g, see Methods for details). The responding light spots could therefore represent neurons with specific orientation selectivity. Thus, the HLC enables a broad-volume analysis of the physiological activity of single neurons in the cerebral cortex.

### 2.3 Functional neuronal imaging of the whole motor cortical area reveals specific patterns of premotor activity representing discrete neuronal assemblies

We next explored the advantages of the wearable instrument for the analysis of brain activity in freely behaving subjects, including movement. For this purpose, the entire motor cortex area of a mouse was exposed on the surface of the brain, such that the HLC could visualize the entire area in both hemispheres simultaneously. Ca^2+^ imaging of CaMK2a-G-CaMP mice was performed using the HLC in the frontal cortical area including the motor cortex of freely moving animals (Supplementary Fig. 8). Different patterns of neuronal activity were elicited depending on different external stimulations of the acoustic and somatosensory modalities. Among those event-related responses, synchronized and slow wavelike activities were observed across broad areas while the mouse was standing still after the stimulation (Supplementary Movie 8).

The pre-motor readiness potential (RP) originally reported as a bereitschaftspotential, is a neuronal premotor activity elicited in the motor cortical area before the initiation of voluntary muscle movement^18^. The RP thought to occur in the higher motor cortex earlier than nerve activity to move the muscle is presumed to be the reflection of the animal’s intention to move the body^19,20^. The premotor activity for an unintended action is categorized as a Type-II RP (no preplanning) while the larger premotor activity for an intended action is distinguished as the Type-I RP (preplanned act); however, it is difficult to analyze premotor activity induced by the random behavior of animals, and the observation of a part of the motor cortex alone cannot reveal the entirety of this premotor activity. Therefore, in the present study, we attempted to measure neuronal activity in the whole motor cortical area of the mouse during voluntary movement to verify that a specific cell assembly expressing the readiness potential might exist.

We developed what we term “restriction motion experiment” to easily detect the activity associated with a specific movement (Fig. 8, Methods section). In this study, a mouse with the HLC on its head was restrained in a plastic tube with its four legs attached to splints set up for monitoring leg movements (Fig. 8a, b). Brain activity preceding the actual execution of exercise, *i.e.*, the readiness potential, is proposed to occur in the motor cortical area^18^, thus we sought to confirm this observation using the HLC on our mouse subjects. Using the movement of a lever as an index, the neuronal activities in bilateral motor cortices were extracted immediately before and after the onset of a specific movement (Fig. 8c, d). First, the locations of neurons activated before and/or after the initiation of motion in the left or right hindleg were either established from raw imaging data by visual judgment of the experimenter (Supplementary Movie 8) or were extracted with a cross-correlation method (Supplementary Fig. 9a). As a result, activity in the M2 area on the contralateral side was higher than that on the ipsilateral side (Supplementary Fig. 9a). Therefore, the right and left hindleg movement accompanied the premotor neuronal activity mainly on the contralateral side of the cortical area, which roughly matches the hindleg position in the secondary motor cortex as estimated previously^21^. However, both these detection methods have intrinsic problems, as visual judgment can easily miss important signals, and screening by cross-correlation can also miss the responses of neurons that do not appear in every trial, but may still be physiologically important. Thus, a more comprehensive quantitative analysis was sought by merging the image data of multiple events (11 left leg kicks, 10 right leg kicks). The maximum FI value during the period (3 sec) before or after the kick was identified for each pixel, and then this value for each pixel was subtracted by the maximum FI value of the same pixel during three 6-sec periods with no leg movement. The subtracted maximum FI value for each pixel was mapped in the right column images of Fig. 8e where the timing of the emergence of each maximum value was color-coded either in green or red depending on whether it appeared before or after the initiation of leg movement, respectively. We further took into account the frequency of these peaks appearing in all kick events, and Fig. 8f shows the time windows before or after the initiation of leg movement during which each peak appeared (see Supplementary Fig. 9b, c for the detailed process). Different levels of color intensity represent the frequency of appearance, and the different colors represent the timing of the maximum peaks. These results also support the conclusion that leg motion is preceded by premotor activity mainly in the contralateral M2 area as a starting point, and as shown here, the activity started within a period of between −2 sec and −1 sec. This timing supports results using the other methods described above, although the distribution does not strictly follow the conventional M1/M2 division, with the activity propagating from the anterior to the posterior poles and from contralateral to bilateral areas broadly. Further qualitative analysis also supported these results (Supplementary Fig. 9d-g), with a cell grouping method based on raw imaging data across the entire motor cortex finding that the cell assembly presenting the pre-motor activities is the main contributor of the lateralized activities both before and after a specific movement (middle graphs of Supplementary Fig. 9d, e). This result suggests that the lateralized activity of the premotor active cell assembly ensures the later lateralized motion. In conclusion, these results indicate that the HLC can visualize specific cell assemblies representing premotor activity in the whole motor cortex (Fig. 8g).

**Figure 8.**
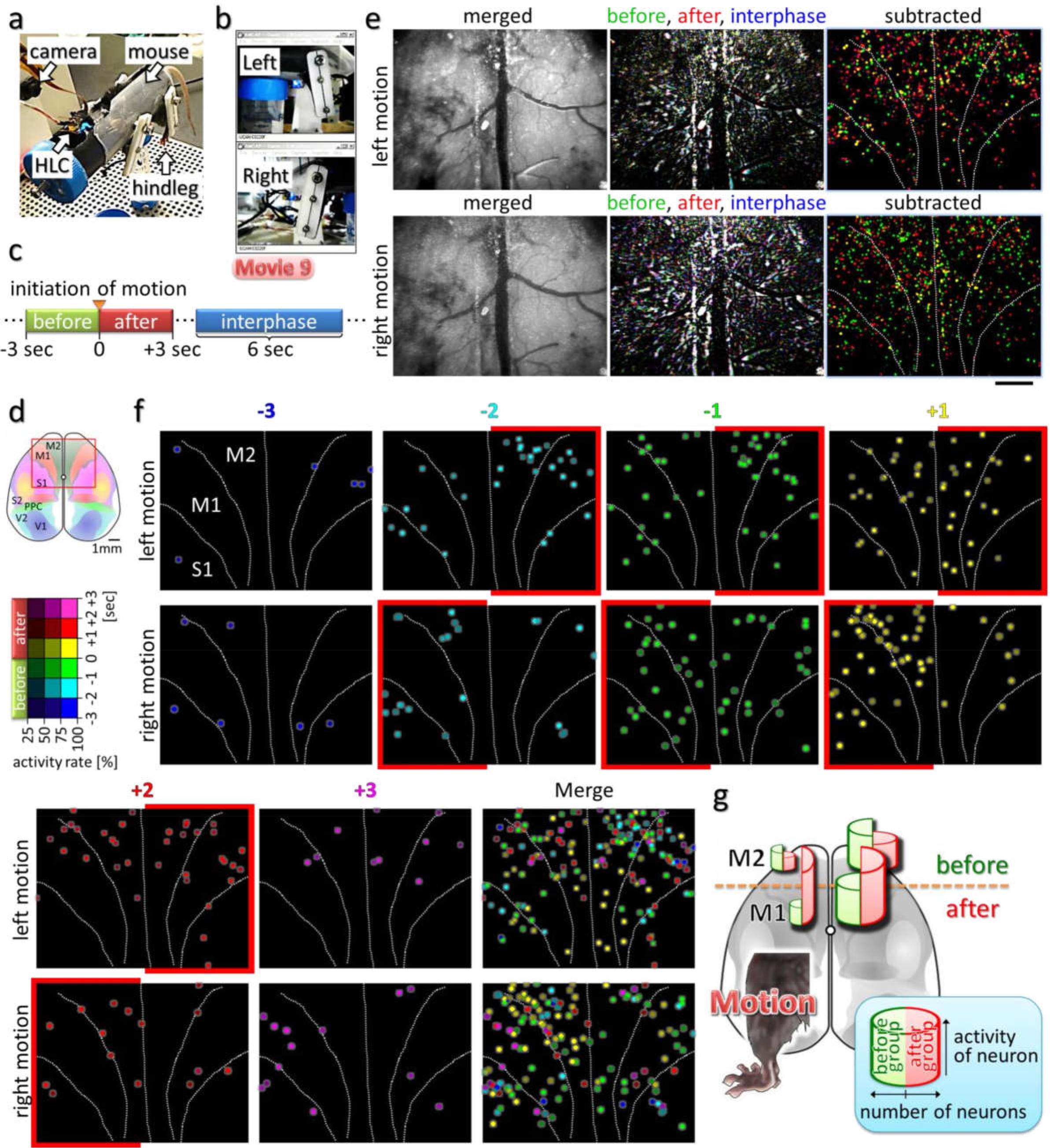
Imaging of the entire motor cortex reveals premotor activity **(a)** The whole view of the restricted-motion experiment apparatus for mice. **(b)** Hindleg movements (kicks) of the mouse were captured by web cameras from both lateral sides of the apparatus. **(c)** Schematic diagram showing classification of the periods during the experiment. The movement state was subdivided into 3-sec durations representing “Before” and “After” the initiation of motion. “Interphase” indicates the 6-sec static state. **(d)** Schematic image showing the HLC imaging position (red square). The Ca^2+^ imaging was performed in awake CaMK2a-G-CaMP7 mice. **(e)** The upper or lower row images show the results of the Ca^2+^ imaging with left or right hindleg kicking, respectively. The left column images represent the fluorescence images in which all raw-imaging data frames in 11 or 10-time left or right kicks during the 6-sec motion state were merged into one frame by maximization. Similarly, imaging data of the “interphase” state in 3-time left or right kicks were merged, and the results are shown in the middle column by blue pseudo-coloring. Also, “before” and “after” states are shown in green and red, respectively. The right column images show the subtracted results, in which green and red represent the [“before” - “interphase”] and [“after” - “interphase”] of the middle column images, respectively. To make visual understanding easier, the light spots of the subtracted images were enlarged and made brighter. **(f)** The subtracted results of (e) for the spots with ΔF/F > 5% were plotted spatiotemporally for each left or right kick (Supplementary Fig. 9c). The brightness of each pseudo-colored dot represents an averaged frequency of appearance of the peak in each trial. **(g)** A schematic diagram summarizing the results in Fig. 8 and Supplementary Fig. 9 for the left hindleg movement. The width and height of the half columns in each hemisphere represent the number and the averaged activity of the “before-” and “after-group” defined in Supplementary Fig. 9d-g. The before-group neurons, which have a peak excitation frequency before the initiation of movement, continue to show lateralized distribution and activity both before and after the motion, while the after-group neurons show less lateralization.

## 3. Discussion

In this study, we describe a wearable imaging system with wide-field, deep-focus optical capabilities that will enable comprehensive physiological analyses of the broad and deep area making up the cerebral cortex. By application of proper image processing algorithm such as NMF, we can also resolve the information of activities of individual cells. In particular, long and continuous (in one day) or longitudinal (over multiple days) observation of the brain in the same individual under free-moving conditions can be performed easily with the HLC. In fact, we observed the light response of the neurons in the visual cortex over multiple days (day1 and 18, or day 11 and 20 after operation). These advantages make the HLC an easy-to-use tool for initial survey analyses prior to using more sophisticated systems for higher resolution studies. To enable better detectability and spatiotemporal resolution, we will need to further improve the optical system and image sensor. In addition, to compensate for the lack of depth information available with HLC imaging, transgenic lines expressing a fluorescent Ca^2+^ indicator could be used to image specific cortical layers or cell types.

Various applications for imaging in the present study revealed the visualization of orientation selectivity encoded in individual excitatory neurons within the whole visual cortex of one hemisphere, and also revealed specific cell assemblies representing activation of the entire bilateral motor cortex during volitional behavior output. The firm attachment of the HLC imaging system to the head of a freely moving mouse enables the stable visualization of the neuronal activities in the same field of view at high reproducibility over many days, and the lightweight property of entire imaging system allows the mouse to move freely with relatively a small amount of stress, also permits the simultaneous use of multiple HLCs (Supplementary Fig. 2, Supplementary Movie 10).

Various volumetric imaging technologies have been developed recently. For example, in multi-photon microscope technology capable of deep observation, a 2P-RAM (2-photon random access mesoscope)^22^ provides a wide field of view with high spatial resolution. However, it is impossible to eliminate the time lag for scanning multiple areas in the large field. A wearable endoscope equipped with a light-field optics^23^ provides 3D imaging with no time lag but with sacrifice of spatial resolution, but it requires a large amount of calculation for the reconstruction of 3D image. On the other hand, a method of 3D imaging by combining a light-field or HiLo (highly inclined and laminated optical sheet) microscopy with a stage system that keeps up with fast motion of an animal was developed^24,25^. Under the necessity to carry out high-speed tracking at high precision in a fixed direction and the limitation in the size of observation field of view and depth, these system have been applied for the whole-brain Ca^2+^ imaging of the freely moving animals with relatively small bodies such as zebrafish larvae. Some of the above methods require expensive and large-scale equipment.

In contrast, among the currently available various wearable optical imaging tools, the HLC is positioned as a unique wide-field, volumetric, and low-invasive imaging device for fluorescence imaging. The HLC in this paper provides a wide field of view and improved detection capability of deep signals. There is no time lag in imaging by the image sensor unlike the galvano scanning system. The laser of the excitation light source provides higher S/N with higher light density than the LED. The spacer apparatus with cranial window can be easily attached to the head with simple surgical operation and enables longitudinal observation over multiple days. Simple mechanisms of the HLC are suitable for commercial production.

Fluorescence change was maintained to be observable as long as 57 days after operation (Supplementary Fig. 7e, f). Judging from the observation by stereo microscopy in Supplementary Fig. 1b, meanwhile, the clarity of the cranial window was kept unchanged for 100 days in the same individual. Therefore, we assume that longitudinal Ca^2+^ imaging is possible for at least 3 months. Because we stopped imaging and monitoring within 3 months, it might still be possible to continue observation beyond this period.

Although only a few animal experiments were conducted to showcase the utility of the microscope in this paper, in the near future, longitudinal imaging using the wearable HLC tracking the same neurons at the same site under free-movement conditions will help to reveal neuronal activity that is specific to various behavioral tasks. Additionally, the simultaneous use of multiple HLCs will facilitate investigation of cognition and behavior through neuronal activity imaging of the global sensorimotor system in animals, especially by a combination of electrophysiology and optogenetical applications.

## Acknowledgements

We thank Charles Yokoyama for helpful comments and for editing the manuscript. The CaMK2a-tTA mouse line was a kind gift of Masako Kawano, Ayaka Bota, and Shigeyoshi Itohara (RIKEN, CBS), and the G-CaMP6/pCAG plasmid was a kind gift of Hisato Maruoka and Toshihiko Hosoya (RIKEN, CBS). Animal care was supported by Yoshie Ito, Megumi Kobayashi, and Kawori Eizumi (RIKEN, CBS). Technical support for the HLC was provided by Hiroyuki Iino (DCT Co., Japan) and Kazushige Ooi (ImageTech Co., Japan), while the 2-photon imaging was supported by Kaori Higuchi of the RIKEN-Olympus Collaboration Center (BOCC). The present study was supported by an internal research budget of RIKEN CBS to H.O. and by the Japan Society for the Promotion of Science (JSPS), KAKENHI grant numbers JP15K21627 and JP17K01996 to T.K.

Construction of the transgenic mouse line was supported by the program for Brain Mapping by Integrated Neurotechnologies for Disease Studies (Brain/MINDS) of the Ministry of Education, Culture, Sports, Science and Technology (MEXT) and the Japan Agency for Medical Research and Development (AMED), and by KAKENHI Grants 15H05723 and 16H06536 from MEXT and the Japan Society for the Promotion of Science (JSPS) to J.N., and by RIKEN through a Grant-in-Aid for Scientific Research for Innovative Areas, namely “Foundation of Synapse and Neurocircuit Pathology” and “Principles of Memory Dynamism Elucidated from a Diversity of Learning Systems” and for Challenging Exploratory Research, and from MEXT, the Human Frontier Science Program, Fujitsu, and Dwango to Y.H., Y.H. is also partly supported by Takeda Pharmaceutical Co. Ltd‥

## Methods

### Development and construction of a wearable imaging system

The wearable HLC imaging system is composed of two major parts, the camera and the spacer (Fig. 1a). The two parts can be separated or combined by using the O-ring as a fastener hooked on their side protrusions. The cylindrical spacer body has a cranial imaging window (object plate) at its base that makes direct contact with the surface of the cortex. The spacer is attached to the mouse head, and the camera is combined with the spacer when imaging is performed. The detachable wearable camera helps with the long-term housing of experimental mice by keeping them free from movement restriction by an electrical wire (Supplementary Movie 1). The camera module (DCT, Co., Japan) has a CMOS (complementary metal-oxide semiconductor) image sensor chip (1/13”, 480 × 640 pixels, pixel size = 1.75 × 1.75 μm; OmniVision Technologies Inc., USA) and a deep-focus optical system. The imaging area is adjustable, and the bottom diameter of the typical cylindrical spacer body is 7.5 ± 0.5 mm. Although the design of spacer body is changeable, typically the height of the spacer body is 9.0 ± 0.5 mm (long spacer in Fig. 5), in this case, the volume of spacer part is approximately 500 mm^3^. The thickness of head cap and gasket is 1.0 mm ± 0.5 mm each, therefore, the total volume of the HLC is approximately 600 mm^3^. The HLC is lightweight (typically 0.9 g) and does not disturb the natural behavior of a small mouse such as individuals from the C57BL/6 line (Supplementary Fig. 2d, e).

The HLC has an LD for the excitation light and an absorption filter for the emission light. The camera was constructed by firmly attaching the micro camera and LD to the head cap with epoxy resin. For green and red fluorescence imaging, the high pass absorption filter set at > 500 or > 520 and > 560 or > 580 nm, respectively (Fujifilm, Co., Japan) was attached in front of the lens, and a blue or green LD (450-460, 488 or 530 nm; OSRAM, Co., Germany) was used, respectively. The LD along with the LSD (Optical Solutions, Co., Japan) and the copper heat sink is driven by the LD driver (Supplementary Fig. 4a, b). The glass object plate (0.525 mm thickness; Matsunami Glass Ind., Ltd., Japan) was firmly attached to the bottom of the spacer body. The head cap and the spacer body were made based on 3D-CAD data by cutting an acrylic plate with a lathe machine.

The image data were transferred to a computer via a USB cable, displayed on a monitor, and saved as an AVI movie file using free software, AmCap (Microsoft, Co., USA). For Ca^2+^ imaging, the image sensor was typically driven at < 30 fps to ensure sufficient sensitivity, stable data transfer and preservation, and real-time presentation on the monitor. The ΔF/F of the mechanical noise when measuring inorganic matter was below the detection limit of 8-bit data. ImageJ (supplied by the National Institutes of Health, USA) was used for all image data analysis. Graphic art works were performed using the free software, DesignSpark Mechanical (3D CAD; computer-aided design, Radiospares Components, Inc.) and GIMP (GNU Image manipulation program, Free Software Foundation, Inc.).

### Verifying the optical specifications of the HLC

The HLC has deep-focus optics and its working distance (WD) can vary by adjusting the lens barrel. The lens position change can also vary with the field of view size at the same time. When the lens barrel is twisted and pulled out for a short WD, the view field becomes small. In contrast, when the lens barrel is twisted and pressed for a long WD, the view field becomes large. If the position of the camera is adjusted properly such that the WD fits the object, the HLC can obtain either a narrow or wide field of view arbitrarily, as shown in Fig. 2a, b. The minimum field of vision is 2.00 × 2.67 mm at 2.70 mm of WD, and here the maximum resolution was 8.33 μm/lp (line pair), 4.17 μm/pixel. The maximum field of vision is 10.0 × 13.3 mm at 15.0 mm of WD, and here the minimum resolution was 41.7 μm/lp, 20.8 μm/pixel.

There is a physical limit to the barrel adjustment range, and if adjusted to an extreme WD value, light aberration cannot be corrected and the peripheral portion of the image is distorted in a pincushion pattern. For instance, with 6.65 × 8.87 mm imaging, the central part of the image can cover 6.65 mm vertically, whereas that of the periphery vertically covers 6.40 mm, which means DTV (television distortion) = (6.65 − 6.40)/6.40 × 100 = 3.91 [%]. Additionally, the central and peripheral parts of the image horizontally cover 8.86 and 8.67 mm, respectively. Therefore, the optical distortion is 2.78 %, calculated as follows: distortion [%] = [(actual half diagonal distance) − (predicted half diagonal distance)]/(predicted half diagonal distance) × 100 = √[(6.65/2)^2^+(8.86/2)^2^] - √[(6.40/2)^2^+(8.67/2)^2^]/√[(6.40/2)^2^+(8.67/2)^2^] × 100. The pincushion distortion will be suitable for observing the convex cortex, but not the barrel distortion (see also right panel in Supplementary Fig. 6b).

Next, the depth of field (DOF) was calculated. DOF is defined by an associated resolution and contrast, and is estimated by a single value calculated from the diffraction limit as a theoretical approximation; however, it is difficult to make a genuine comparison because many imaging lenses are not diffraction limited. Therefore, the only way to truly determine DOF is to use a test target. Normally, even if a lens has infinite focus theoretically, the spatial resolution of the image sensor (density of the photo-diode pixel array) is limiting for DOF. Firstly, the line spread function (LSF) of the HLC in the case of 6.65 × 8.87 mm imaging (resolution; 27.7 μm/lp, 13.9 μm/pixel) was analyzed by using a line chart (Fig. 2c, d), and then based on the results of the LSF, the square wave response function (SWRF) was calculated as follows:

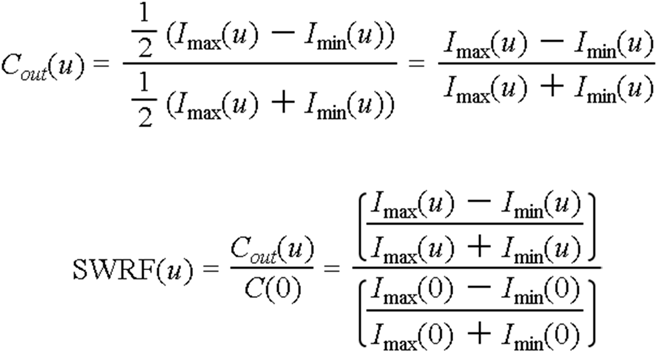

*C_out_*(*u*) is the output contrast of rectangular wave pattern at spatial frequency *u*. *I_max_* and *I_min_* are the values obtained by converting the intensity into the dose at each square wave of the LSF. As a result, the SWRF is 24.7 [lp] at a cut-off point of 0.03, as generally used by many optical manufacturers (Fig. 2e). That SWRF value corresponds to the position at 600 pixels in LSF (Fig. 2d). In the line chart, width is 0.912 mm. Therefore, DOF is 24.7 × 0.912 × √3/2 = 19.5 [mm].

Finally, to calculate an effective spatial frequency at the distorted edge of the imaging field, the SWRF (a rectangular wave response function) was corrected to the modulation transfer function (MTF; sine wave response function), calculated with the correction via Coltman’s formula^26^. MTF is a measure of an imaging lens’s ability to transfer contrast from the object plane to the image plane at a specific resolution, and is expressed with respect to image resolution (lp/mm) and contrast (%). Typically, as resolution increases, contrast decreases until a cut-off point, at which the image becomes irresolvable and grey. The formula is shown below.

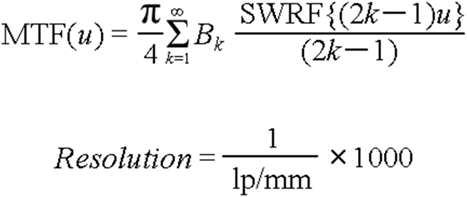

If the total number of prime numbers in (2*k* − 1) is *m* and the number of types of prime numbers is *n*, *Bk* = 0 when *m* > *n*, and *Bk* = (−1)*^n^*(−1) ^*k* −1^ when *m* = *n*. For the calculation of MTF, up to the fourth term of the expansion formula was used according to a previous verification^27^. As a result, MTF resulted in an effective spatial frequency of 15.5 lp/mm where MTF = 0.03 (Fig. 2f), producing an effective spatial resolution of 64.5 μm/lp. Therefore, the estimated minimum effective spatial resolution was 64.5 μm/lp at the distorted edge of the imaging field, which is 2.3 times that of 27.7 μm/lp at the center of the visual field in the case of 6.65 × 8.87 mm imaging (resolution; 27.7 μm/lp, 13.9 μm/pixel). According to the above results, although actual measurement values will fluctuate depending on the measuring environment and optical distortion, a discriminable minimum light spot is presumed as approximately two times blurred at the edge of the field of view due to natural optical aberration.

### Quantitative imaging analysis using fluorescent beads

After craniotomy, fluorescent beads (F21010, green fluorescent FluoSpheres, polystyrene; Thermo Fisher Scientific Inc., USA) were implanted into the cortex of the anesthetized mice at different depths by using a needle and micromanipulator (Fig. 3). The bead diameter of 15 μm was similar to the general cell size. The stereomicroscope has a general objective focus lens, whereas the HLC has a deep-focus lens. In Fig. 3, the field of view of the HLC is 3.80 × 5.06 mm, 1 pixel = 7.917 μm. A general objective lens can handle a bright image because the F value is smaller than the deep-focus lens, and a sharper image was obtained by focus imaging in Fig. 3b than in Fig. 3c; however, the DOF was shallow. In contrast, with the HLC, the fluorescence derived from every bead can be detected even at an 800 μm depth, suggesting that the detectability of fluorescence by the deep-focus lens is apparently higher than that for a conventional focus lens. In the left 2 images of Fig. 3d, the position of the upper beads at 400 μm depth does not match exactly between the conventional microscope and the HLC. This is likely due to differences in optical distortion. Regarding the small 10 images to the right column in Fig. 3d, although these images are shown brightly for the sake of convenience in aligning and distinguishing shapes, the measurement was actually performed based on the raw data, with the quantitative results shown as a graph in Fig. 3e, f. In the microscope image, the FI and shape of each bead at the same depth are almost the same (left graph in Fig. 3e), supporting the accuracy of the experimental system. Thus, the deeper the position of the beads, the lower the FI becomes in the image taken by the conventional stereomicroscope, until the FI is reduced almost to background levels at 800 μm. In contrast, the decrease in FI with depth is mild for the HLC imaging, and a stronger fluorescent signal than for the conventional microscope was detected even at a depth of 800 μm. Compared at the same depth, the bead’s FI level and shape captured by the HLC are slightly variable. This can be reasonably explained by distribution differences in the excitation light (see also the image in Fig. 2h) and optical distortion (as mentioned above in Methods section for Fig. 2a). In fact, from the center of the field of view to the periphery, the intensity of the excitation light decreases and the size increases. It is possible to correct the above-mentioned variability mathematically, if necessary, in actual functional brain cellular imaging. Concerning the result of Fig. 3f, the deeper the position of the bead, the larger it becomes relative to its actual size in the conventional microscope images. In contrast, image sizes with the HLC are generally constant relative to depth. Specifically, the enlargement ratio at 50% FI with the stereomicroscope image actual bead size is 4.10 (at 100 μm), 14.6 (at 400 μm), and 17.6 (at compared to 4.75 (at 100 μm), 5.80 (at 200 μm), and 6.86 (at 800 μm) for the HLC imaging. Thus, both the decline in signal detection ability and the expansion of the outline seem to be less when using the HLC due to the deep-focus optics.

### Ca^2+^ imaging in 3D culture cells

Cell cultures in 2D and 3D, gene transfection, and drug administration were performed according to previous reports^9,28^. Hela cells, derived from a human cervical cancer, were transfected with the G-CaMP6 gene^12^ under 2D conditions.

After transfection, cells were embedded in the collagen gel as an extracellular matrix to imitate brain structure as a brain phantom. Hela cells are activated by histamine through the histamine H_1_-receptor by producing intracellular Ca^2+^ increases^29^. Thus, when a histamine solution (final 5 μM) was applied to the dish, rising Ca^2+^ increases were detected by the HLC at 30 fps imaging (Supplementary Fig. 6a, Supplementary Movie 4). In contrast, the increase was abolished by 5 mM EGTA (Fig. 4b).

### Animal studies

All procedures involving animals conformed to the animal care and experimentation guidelines of the RIKEN Animal Experiments Committee and Genetic Recombinant Experiment Safety Committee. C57BL/6J (SLC Co., Japan), CaMK2a-tTA (Jackson 3010)^30^, TRE-G-CaMP7-2A-DsRed2^31^, and Thy1-G-CaMP7-2A-DsRed2 (Thy1-G-CaMP7) ^32,33^ mice, aged 6-12 months, were used for the *in vivo* experiments (see details of surgical operation in Supplementary Fig. 1).

### Cell detection and signal extraction

In Fig. 6c, imaging data analysis was performed using custom codes in LabVIEW (National Instruments) and Matlab (Mathworks). Brain imaging movies were processed in the following steps to detect cell bodies and extract their temporal signals. First, for each pixel of the image, a variance value was calculated using the time signal of that pixel. Thus a variance map was obtained for each movie. This variance map has peaks and valleys of various heights, corresponding to the location cell bodies. This variance map was then cut horizontally at different heights to obtain 100-200 slices, depending on the data. Each of these slices showed presence of segmented ROIs. Cells with weaker activation were visible in slices cut in higher depth from the top, while highly activated cells were found in slices in lower depths. All ROIs with area in the range of 30-150 squire microns were gathered to form a preliminary set of cell-like ROIs. For each ROI, a 2-pixel-wide Gaussian filter was used to smooth the spatial distribution of the ROI. However, as in many cases same cells were detected multiple times in varying depths, temporal correlations among these ROIs were checked. ROIs having temporal correlation value >0.9 and some degree of spatial correlation were considered as same cells, and they were added to form one single ROI. By performing this task recursively, we obtained a second set of ROIs representing the cell bodies in the movie.

In the next step, we calculated temporal signals of these ROIs from the movie by taking weighted average of temporal signal of all the pixels in a specific ROI. As the intensity of the movie pixels varied due to position of pixels, ROIs has various levels of baseline intensity signals. Furthermore, in case of weakly activated cell bodies, actual calcium transient signal was not clearly visible due to temporally fluctuating baseline. Therefore, to eliminate baseline effect, we used a NMF (Non-negative Matrix Factorization) algorithm^14^ to extract actual activation of the cells while separating the spatio-temporal baseline of the image simultaneously by solving the following problem:

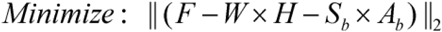

*F*: orginal image
*W*: spatial components corresponding to cells
*H*: temporal components corresponding to cells
*S_b_*: Spatial baseline
*A_b_*: Temporal baseline

With the application of the above mentioned algorithm, baseline removal and separation of temporal signals corrupted by spatial overlap of cells was achieved.

### Perceptual and behavioral tests

In Fig. 7, awake mice received either light stimulation or an 8-direction drifting grating with blue light stimulation using a light emitting diode (LED, peak wave length 470 nm; Stanley Electric Co., Ltd., Japan) driven by a function generator, or video stimulation using a computer monitor. C57BL/6 mice have high photosensitivity against blue light judging from their pupillary light reflex^34,35^. Thus, to increase the stability of repeated visual inputs to the mouse, the head of the mouse was fixed with a stereotactic instrument while its body was allowed to move freely (upper left image in Fig. 7a), so that the left eye was given the visual stimuli while the right eye was masked. All experiments were done in a dark room, and the imaging was started after 15-minute habituation. In Fig. 7a, the upper middle schematic diagram for the rodent brain was drawn with partial modification by reconstructing the transverse sections of the brain atlas^36,37^ (Supplementary Fig. 8a).

A single flashlight stimulation (rectangular pulse 0.1 sec) or 10-times repeated flashlight stimulation (rectangular pulse trains of 0.5 sec, 20 Hz, duration 25 ms, interval 25 ms) were applied to the mouse. Similarly, the 8-direction drifting grating (each drifting duration 3 sec, interval 6 sec, > 0.5 cycle/degree^38^) was presented to the mouse. Ca^2+^ imaging was then performed using the HLC on the CaMK2a-tTA × TRE-G-CaMP7-2A-DsRed2 mice (CaMK2a-G-CaMP7) over the right visual cortical area. ROIs of 4 pixels were selected by visual judgment, and changes in FI were measured from the raw imaging data. Fig. 7b represents the results of 12 ROIs of 2 mice in 6 trials, while Fig. 7c shows the results of 7 ROIs of 2 mice in 3 trials. All ROI positions are shown in the lower column images of Fig. 7a. In Fig. 7d, the left and right graphs plot the raw values obtained from 15 ROIs of 2 mice in 3 trials and 12 ROIs of 2 mice in 1 trial, respectively. ROIs of one mouse (ID: #2) among 2 mice is exemplified in Fig. 7e. In Fig. 7f, Data of 4 opposite angles ([0 and 180 ^o^], [45 and 225 ^o^], [90 and 270 ^o^], and [135 and 315 ^o^]) during the 3-second stimulation were maximized, and data of right angles were subtracted as below, ΔF [0 and 180 ^o^] = FI ([0 and 180 ^o^] − [90 and 270 ^o^]), ΔF [45 and 225 ^o^] = FI ([45 and 225 ^o^] − [135 and 315 ^o^]), ΔF [90 and 270 ^o^] = FI ([90 and 270 ^o^] − [0 and 180 ^o^]), ΔF [135 and 315 ^o^] = FI ([135 and 315 ^o^] − [45 and 225 ^o^]). Finally, these four different ΔF’s were assigned with different 4 colors (Green, Blue, Magenta, Yellow), and each ROI was given a pseudo-color by weighed superimposition of these 4 colors to represent the orientation preference of each ROI (Fig. 7f). Among these ROIs, those with top 3.5 % of pseudo-color intensity are presented in Fig. 7g.

Regarding deconvolution (upper right image in Fig. 7a), diffraction point spread functions and iterative deconvolutions were calculated according to Dougherty’s algorithm^39^. In the present study, FI measurements were always extracted from raw data, and not from the deconvoluted data.

For the maximum points extraction (Supplementary Fig. 3), light points were selected and counted from the image after subtraction of background with the parameter of a noise tolerance of 10 according to the ImageJ algorithm constructed by Michael Schmid (NIH, USA).

For the “restriction motion experiment” represented in Fig. 8, an apparatus was made to analyze neuronal activity in the mice derived from specific voluntary movement initiation. A cylindrical restraint tube was filled with urethane foam resin to fit the body shape of the mouse, and its external wall was painted with black ink. Hence, the body of the mouse is held firmly when it is inserted in the tube, and other parts of the body except the legs do not move as much, thus mildly restricting the mouse motion and sensation. Before beginning the measurements, the mouse was placed in the cylindrical restraint and their legs protruding from the tube were attached to splints, which functioned as body-worn foot levers. The splints could move according to the leg motions or could be made immovable individually by the fixing of bolts. The motion of the splint was followed by the movement of a line drawn outside the splint and captured by a web camera at 30 fps from both lateral sides. Then, Ca^2+^ imaging was performed after 15-minute habituation in 4 mice (ID: #3-6) using an HLC attached to the spacer protruding from the upper part of the restraint tube, at 20 fps (Fig. 8) or 10 fps (Supplementary Fig. 9a, Supplementary Movie 9).

A series of analyses comprising a “comprehensive quantitative analysis” (Fig. 8e, f, Supplementary Fig. 9b, c) and an “overall qualitative analysis” (Supplementary Fig. 9d-g) were also conducted as described in the respective figure legends. The maximization of FI in the imaging data, described as a maximum intensity projection in Supplementary Fig. 7, removed the basal random noise similarly to an averaging process, but without losing the information of relatively rare but important events.

## Data availability

All relevant data are available from the authors.

## Supplementary information

**Supplementary Fig. 1.**
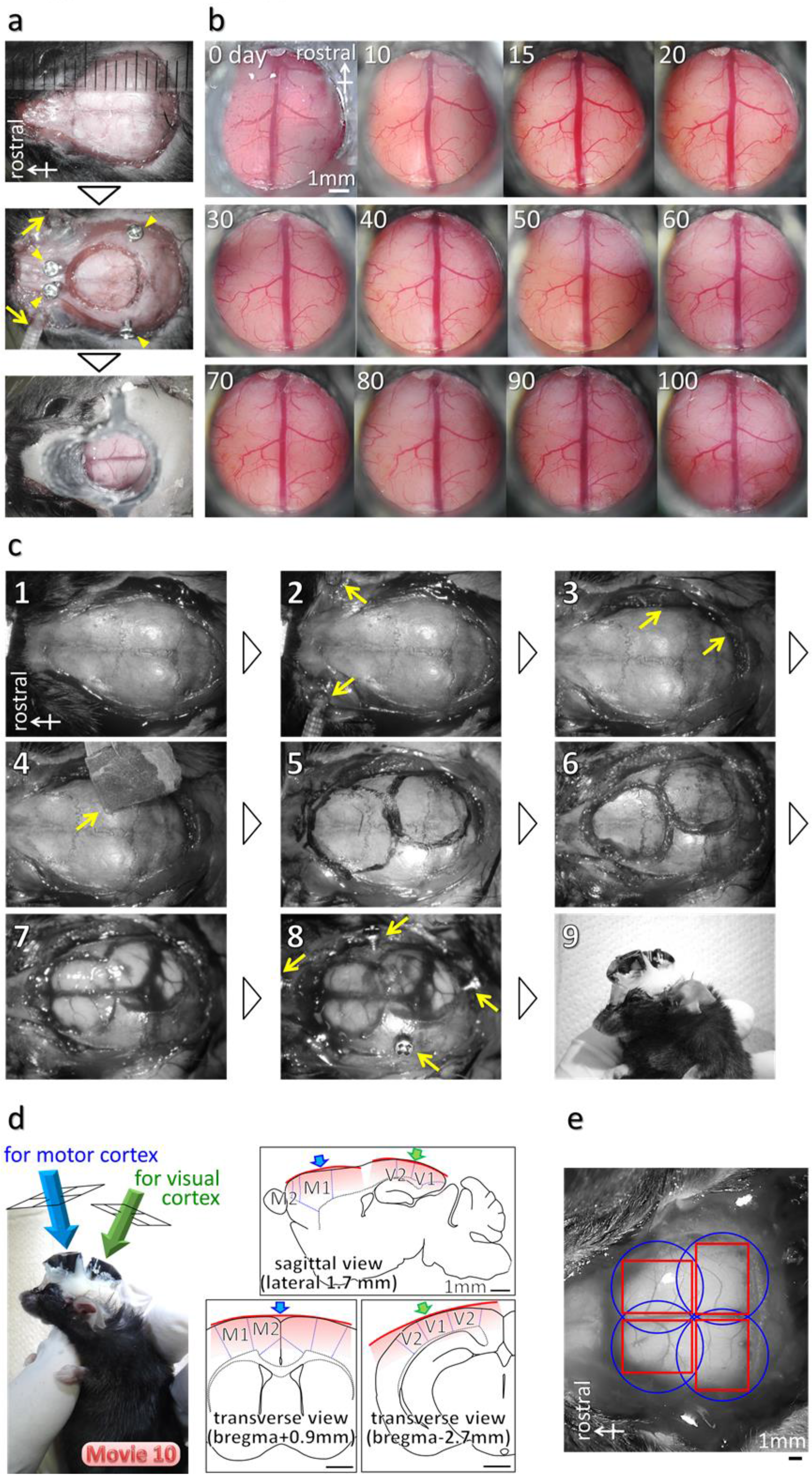
The HLC allows long-term observation of the same brain area in the same mouse. **(a)** Typical surgical operation and the process of the applying a single use of the HLC. The mouse was anesthetized with 2,2,2-tribromoethanol (125-250 mg/kg body weight^40^) or a combination anesthetic consisting of 0.3 mg/kg medetomidine, 4.0 mg/kg midazolam and 5.0 mg/kg butorphanol^41^, and was mounted on a stereotaxic instrument (Narishige Co., Japan). The head skin was scalped (upper panel). After craniotomy using a dental drill, the screws were inserted at bone positions around the hole to anchor the dental cement (yellow arrowheads in the middle panel). Eyelids were clamped with small forceps for protection (yellow arrows). The dura mater of the mouse was removed by using a hand-made sharpened tungsten needle gently and carefully. And then, the cylindrical spacer with a cranial window for imaging was attached to the mouse head by using dental cement (UNIFAST, GC, Co, Japan) (lower panel). **(b)** Images show the temporal serial observation of the same mouse using a stereo microscope. Some part of the brain surface causes bleeding just after removing the dura although the extent of bleeding depends on operational skill. Bleeding was promptly stopped and the microvessels recovered in 4-5 days. Thereafter, the observation surface was kept clean for more than 100 days. During this long-term housing, neuronal activity could be visualized using the HLC and conventional 2-photon microscopy through the cranial window of the spacer apparatus (Supplementary Fig. 7). **(c)** For multi-point imaging using two HLCs, surgical operation and the processes are shown. The detailed order of each process is described below; 1) Scalp removal. 2) Eyelid clamping using a small forceps for protection (yellow arrows). 3) Peeling part of the muscles of temporal and occipital regions (yellow arrows). 4) Removing the periosteum with water-resistant sandpaper (yellow arrow) (*e.g.* #600 waterproof paper file). 5) Marking the craniotomy position. 6) Drilling the skull by using a dental drill. 7) Removing the skull (optional: removing the dura mater). 8) Anchoring the screws (yellow arrows) (*e.g.* #0 pan head, M1.0 × 2.0 mm). 9) Attaching the cranial window parts with dental cement. Two cylindrical spacers with cranial windows were put on the cortex by using a precision manipulator to firmly support their positions, and then the spacers were fixed to the skull with dental cement. Besides, for behavioral testing under bright environments, black liquid rubber may be applied around the spacer and the dental cement for light interception. **(d)** Example of installation of duplicate spacers for visual-motor imaging in the freely moving CaMK2a-G-CaMP7 mouse (Supplementary Movie 10). Blue or green arrows indicate the direction of observation from the HLC for imaging including the whole motor area of bilateral cortex or the whole hemispheric visual area, respectively (left picture). The right panels show the directions of observation and the thick and convex area of imaging by the HLC (red line and graded red area, see also right image in Supplementary Fig. 6b) both in the sagittal (upper panel) and the cross (lower panels) sections. Use of multiple cameras allows the broad area imaging of the curved cortex at cellular resolution (>4.17 μm/pixel). **(e)** Schematic image of the application of four HLCs. Blue circle and red square indicate the position of the spacer and the imaging area, respectively. The quadra imaging achieves simultaneous imaging of the entire cerebral cortex. As mentioned in Fig. 2a, one HLC can capture a wide field of view of 10.0 × 13.3 mm. This size is almost equal to one brain of the mouse. However, in that case, the resolution is lowered to 20.8 μm/pixel, and the sight from a single direction distorts the image of the spherical brain near its periphery. With multi-point thick imaging, it is possible to image at a high resolution with low distortion along the curvature of the cortex.

**Supplementary Fig. 2.**
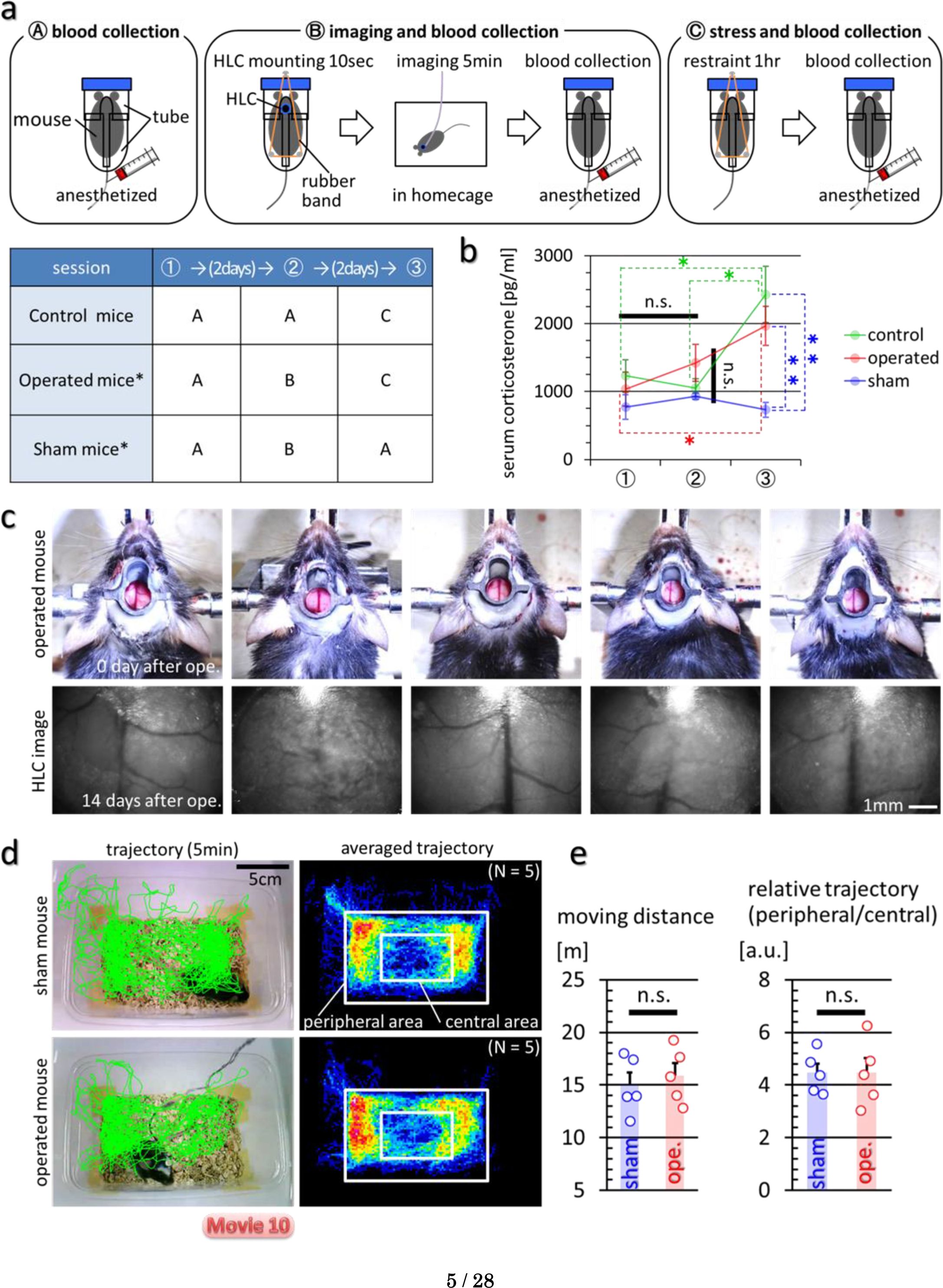
The imaging of neuronal activities by the HLC attached to the head of freely moving mice causes no significant increase of stress, and does not affect the locomotor activity and the behavioral pattern. Blood-based and behavioral assessments of stress caused by wearing the HLC were performed. **(a)** Corticosterone produced in response to stress was measured. Schematic images of experimental procedures are shown in A-C (upper images). These 3 different experiments were conducted in 3 sessions every 3 days for 3 groups of mice (each N = 5, total 15 mice) (lower table). 8-10 months old C57BL/6JJmsSlc male mice were purchased (Japan SLC, Inc., Japan). A group of mice with an asterisk were given the 5 minutes daily handling for 1 week prior to the experimental session. “Operated mice” were attached with the spacer apparatus with cranial window on his frontal cortical area by surgical operation performed 2 weeks before this experiment. “Sham mice” are the wild-type mice which underwent the same treatment as in the procedure B but were not mounted with the HLC. **(b)** The result of ELISA assay of the serum corticosterone level is shown. All procedures were performed according to the manufacturer’s protocol (Corticosterone ELISA Kit, Cayman Chemical Company, USA). Error bar is a standard error of the mean. T-test was performed after F-test for judging significance. Single asterisks mean there is the significance of p < 0.05 between session 1 and 3 (p = 0.0271), session 2 and 3 (p = 0.0137) in control mice, and between session 1 and 3 (p = 0.0213) in operated mice. Double asterisks mean there are significance of p < 0.01 between control and sham mice (p = 0.0067), operated and sham mice (p = 0.0049) in session 3. There are no significance (n.s.) between session 1 and 2 in all mice group, and between control, operated and sham mice in session 2. These results indicate that the HLC mounting and imaging process causes no significant increase in stress in the mice. **(c)** These photos show the individual operated mice and their frontal cortical image taken by the HLC under the freely moving condition (“ope.” means the surgical operation). **(d)** The locomotor activity in the homecage from which the lid was removed was examined. Left column images indicate the examples of the moving trajectory for 5 minutes by a sham and an operated mouse (Supplementary Movie 10). Right column images indicate an averaged trajectory of 5 sham and operated mice. The averaged trajectories were binned with 5 × 5 pixels, and shown with pseudo-color. The bottom of the homecage was subdivided into the central and peripheral areas as indicated in the figures. **(e)** The numerical analyses of (d) are shown in the graphs. Error bar is a standard error of the mean. Each circle indicates the value of each individual. There is no significance (n.s.) between the sham and operated mice in the moving distance or the distribution of the trajectory. These results suggest that HLC mounting and imaging process does not affect the locomotor activity and the behavioral pattern.

**Supplementary Fig. 3.**
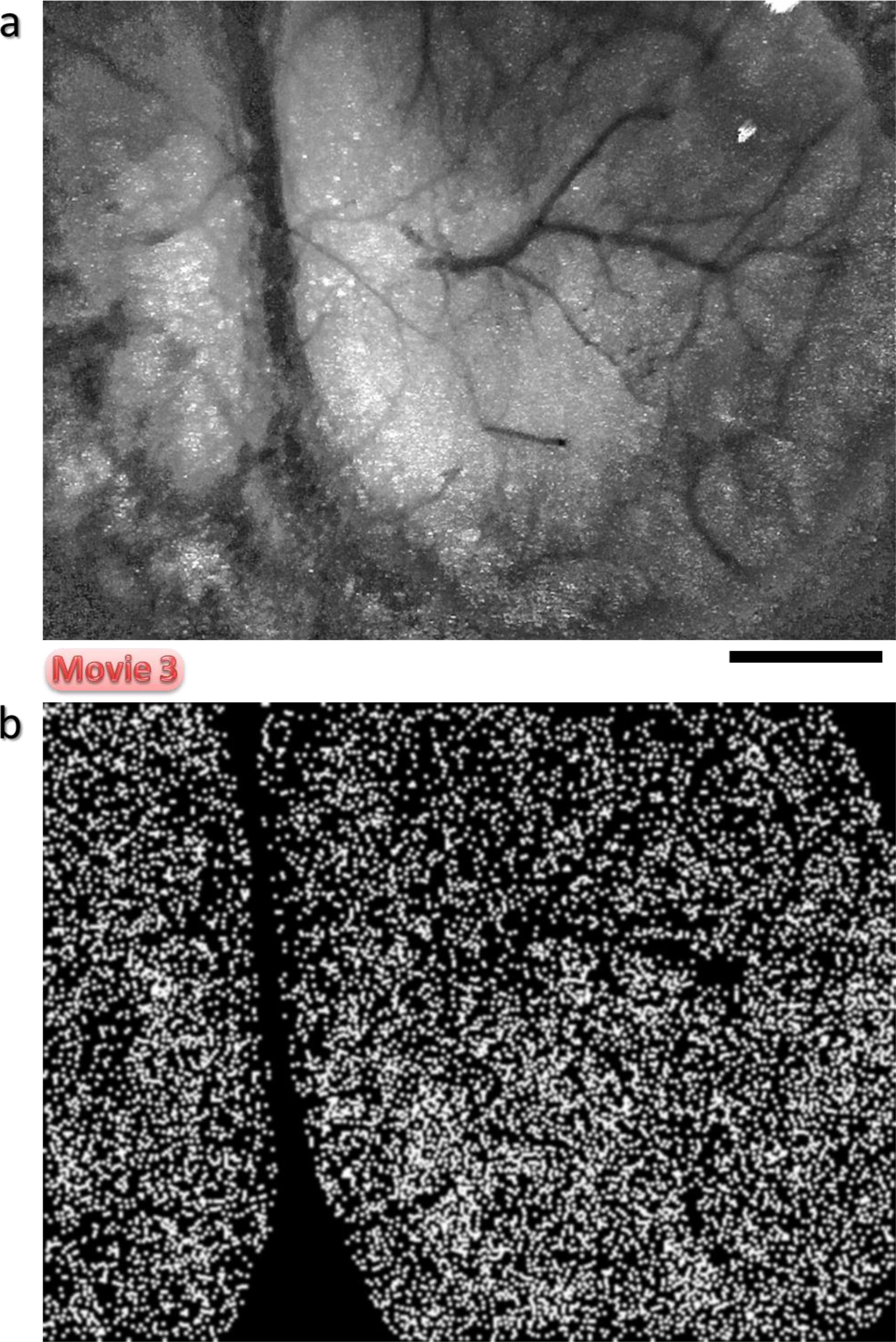
Automated selection and counting of discriminable blinking light spots. **(a, b)** Ca^2+^ imaging was performed by the HLC on the visual cortex of the CaMK2a-G-CaMP7 transgenic mouse (Fig. 7). In order to estimate how many excitatory neurons can be detected and discriminated in the occipital area including the visual cortex by the HLC, the merged image was analyzed by counting the number of peak fluorescence spots (see for detail Methods section). 8 raw movies for 1 min were taken by the HLC and merged by maximization. A representative 10 seconds of the merged movie is shown in Supplementary Movie 3, and the results from where all frames were merged by maximization is shown in (a). Bar indicates 1 mm. The maximum points of the fluorescence spots of (b) are shown in (a). 9352 light spots could be discriminated within a single field of view by one HLC (4.25 × 5.66 mm, 30.7 × 10^4^ pixels, 8.8 μm/pixel, 1-6 pixel/spot in (a), 5 pixel/dot in (b)).

**Supplementary Fig. 4.**
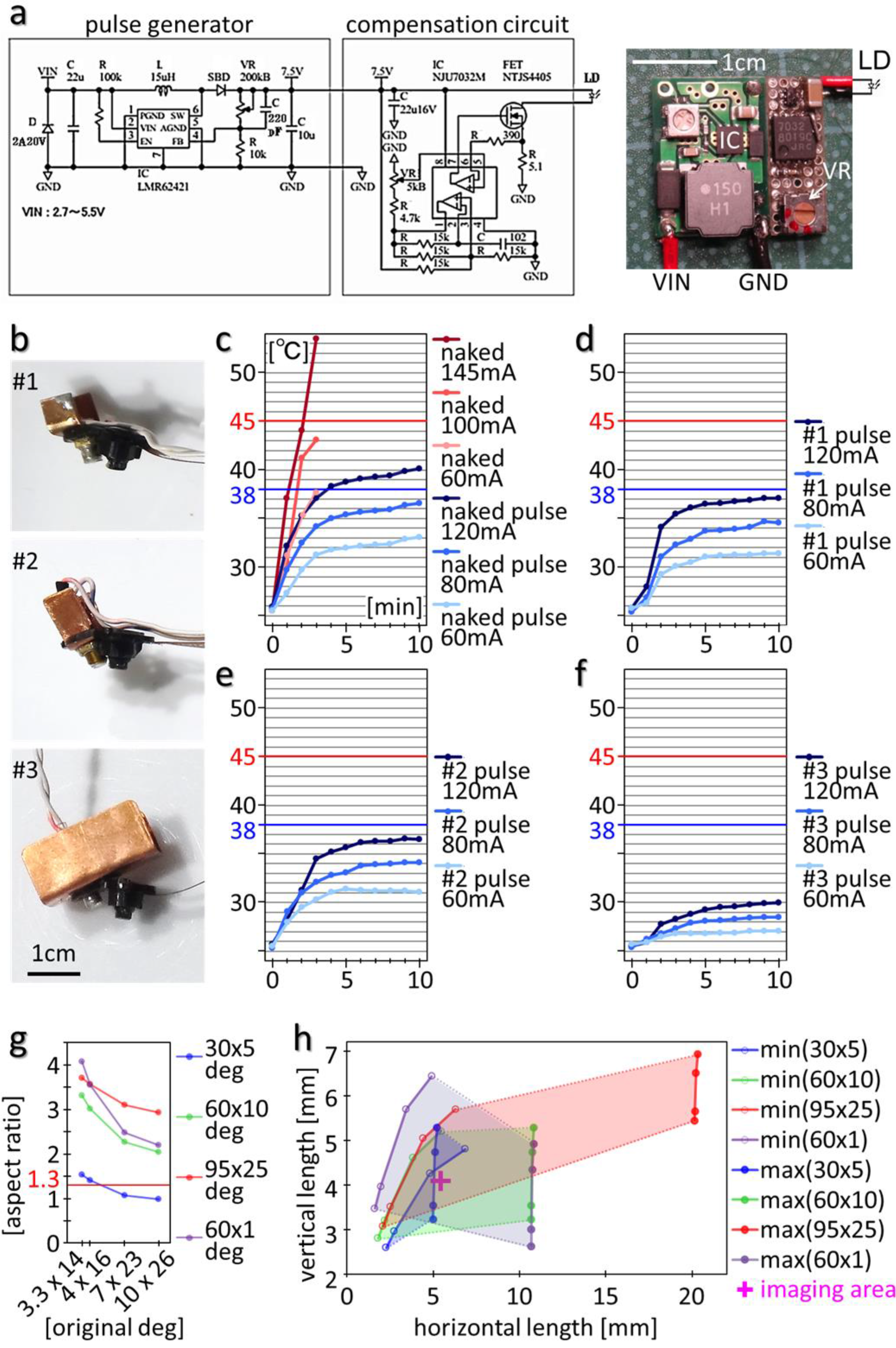
Components: a pulse generator to drive a laser diode (LD), an external heat sink structure to suppress the temperature rise of LD, and a light shaping diffuser (LSD). **(a)** The LD driver and its circuit diagram. To prevent a temperature rise of the LD due to continuous illumination, a LD driver (ImageTech, Co., Japan) was regulated by a pulse generator which turns the LD on and off at a high-speed. In the circuit diagram, the left box shows an oscillator and the right box shows a compensation circuit. 32 kHz pulse is stably supplied to the LD. Since a source voltage allows 2.7-5.5 V input, the LD driver can be driven by an external power supply unit or also directly connected to the USB power supplying wire, which contributes to the miniaturization and simplification of the entire system. A CMOS operational amplifier was used in the compensation circuit for a constant current. MOS-FET (metal-oxide-semiconductor field-effect transistor) was used for an output enhancement as a switching element. Output power for LD can be controlled by a variable resister (potentiometer) (VR, white arrow in right image). Therefore, it is possible to irradiate excitation light with a preferable output according to the observation target during the imaging. VIN, input voltage; GND, ground; C, capacitor; R, resistor; L, inductor; D, diode. **(b)** A heat sink of variable size was attached to the LD of the HLC. #1/ #2/ #3 shows 146.7/ 194.7/ 889.2 mm^2^ heat sinks, respectively. **(c)** The temperature change of the LD (PL450B, OSRAM, Inc., Germany, threshold current 30 mA, maximum optical output power 100 mW, threshold current 30 mA, operating current <145 mA) was examined. The temperature of the metal housing of the LD attached to the heat sinks was measured using a thermocoupled probe. The room temperature was 25.5 °C. Blue or red horizontal lines in each graph indicates 38 or 45 °C respectively, as a typical mouse’s body temperature or as a presumptive limit temperature at which irreversible damage is caused to the cell. When a constant current of 60, 100, or 145 mA was applied to the “naked” LD (no heat sink), its temperature rapidly exceeded 38 °C (c). In contrast, when a pulsed constant current (32 kHz) of 60, 80, 120 mA was applied using a pulse generator, the temperature of the naked LD increased more slowly, finally reaching a plateau level after 10 minutes, thus showing the effectiveness of the pulse drive for stable lighting by the LD and the prevention of a rapid temperature rise. **(d-f)** The temperature of the LD with the heat sink was measured. These results indicate that the cooling efficiency by the heat sink is higher as its surface area increases. The temperature of the LD with every type of heat sink reached a plateau without exceeding 38 °C, even if they were driven at 120 mA. Therefore, the heat sink structure is effective for cooling the LD. Also, the time of LD with the #1 heat sink temperature to reach 95 C of its maximum temperature was more than twice as fast as that of the naked LD. From a practical viewpoint, we decided to use the #1 sink in our *in vivo* experiments. It is necessary to change the current value of laser diode (LD) according to the observation target. The experimenter needs to dim the LD while observing the fluorescence of the target during the experiment. As far as using the HLC (image sensor is OV7690) with the CaMK2a-G-CaMP7 mouse, it was mainly used at under 60-80 mA in most cases. At this situation, the temperature is 33.0 − 36.5 °C and it is below the body temperature of the mouse, so we do not need the heat sink (c). Since the LD is not in contact with the mouse and it is installed away from the mouse, even if it reaches 40 °C at 120 mA, it is considered that there is practically no problem if lighting is limited to short time. On the other hand, it is considered preferable to use a heat sink if it is expected to be used for longer period at approximately 90 mA or more. In this condition, LD is expected to exceed the mouse body temperature (depending on the room temperature). The heat sink suppresses the temperature rise above the room temperature, and shortens the time to reach the plateau, so it stabilizes the laser light quickly. Therefore, the experimenter can shorten the standby time associated with turning on and off the LD. In summary, if the experimenter emphasizes weight reduction of the HLC, it is not necessary to attach the heat sink. While, if the experimenter wishes to give priority to shortening the standby time, it is effective to attach the heat sink. **(g, h)** The LD is used as the excitation light source of the HLC. The laser light from LD has a flat shape (lower leftmost panel in Fig. 2h). Therefore, the irradiation beam was processed by using a light shaping diffuser (LSD). In the right panel of Fig. 2g, the LD was tilted 30 degrees from the light axis of the camera and the paper was illuminated from a distance of 8 mm, which is the same condition as actually used in the HLC. As a control, we used a white LED that projected a round light bundle. The projected images by the white LED with a circle lens or LD (PLT5 488, OSRAM, Inc., Germany) were taken in the dark by a digital camera (COOLPIX P7100, Nikkon, Inc., Japan) with ISO100, exposure time 1 second, and fixed focus. The oscillation threshold of the LD, PLT5 488, is 30 mA, and the maximum limit current is 150 mA. According to the specification, the original beam divergence changes in the range of 4 × 16 − 7 × 23 (at 75 mA) - 10 × 26 degrees (deg) depending on the amount of applied current. Since the LD is usually driven at 40-80 mA for *in vivo* imaging, the distribution of LD at 40 mA was first analyzed. In the upper line of Fig. 2h, the projected circle light by the white LED was processed to a horizontal ellipse shape with various LSDs; original, 30 × 5 deg, 60 × 10 deg, 95 × 25 deg, 60 × 1 deg. Similarly, the figures in the lower line show that the projected vertical elliptical light from LD was processed to horizontal ellipse shape. In the result of the light projection using the 60 × 10 deg LSD, the distribution of a yellow-colored area with a half-value intensity was within an inner red square which corresponds to the typical view field of the HLC (4.0 × 5.3 mm). Therefore, we thought that the light distribution processed by the 60 × 10 deg LSD most closely matches the region of the imaging field of the HLC. Continuously, the aspect ratios of the diffusion angle of the projected beam by the various LSDs were calculated depending on the original divergences of the LD at different currents (g). And finally, the various projected ranges by the various diffusion angles were simulated (h). The graph in (g) indicates the aspect ratio of the vertical and horizontal projected angle in each diffusion angle LSD (indicated by different colored lines) plotted against various divergences of the original LD beam at different currents. X-axis values are plotted according to the horizontal degree of the original LD beam. The aspect ratio of the imaging area of the CMOS, *i.e.* 640 × 480 pixels, is 1.33 (led line). When the LSD is applied, the diffusion angle is approximated by the following formula: √[(light source divergence angle)^2^ + (LSD diffusion angle)^2^], and the diffusion range is approximated by the following formula: (the distance to irradiated surface) × 2 tanθ, whereθ, is the half of the diffusion angle. In this experiment, the light source position was tilted by 30 deg horizontally. Therefore, the distance to the irradiated surface was 8 mm × 2/√(3), and the horizontal diffusion range was also corrected by multiplying 2/√(3). From the measurement result based on the projected original image in Fig. 2h, the original beam divergence of LD was 3.3 × 14 deg at 40 mA. In the case of 3.3 × 14 deg, the shape tendency calculated by the corrected angle at various aspect ratios of the LSD coincides with the actual distribution in Fig. 2h. As a result, a 30 × 5 deg LSD is the most preferable and 60 × 10 deg LSD is the secondarily preferable for all current ranges of the LD. The graph in (h) indicates the various projected ranges (vertical, horizontal length) by the various diffusion angles of the LSD (indicated by different colored lines) at the minimum or maximum current of LD. The diffusion angle increases or decreases according to an increase or decrease of the current value. The minimum projected ranges with various LSDs by the minimum current were calculated based on the yellow-colored area of projected image of Fig. 2h, and are shown as “min (LSD variation)” on the right side of the graph. Similarly, the maximum projected ranges were estimated based on 3.3 × 14, 4 × 16, 7 × 23, 10 × 26 deg as a maximum original divergence, and are shown as “max (LSD variation)”. The four dots in the data for each LSD show the results in order of the current value. The colored area for each type of LSD indicates the projected range that can change from the min value to the max value. “Imaging area” (magenta cross in the graph) indicates the typical HLC imaging area (4.0 × 5.3 mm). As a result, all tested LSDs except the 30 × 5 deg LSD can cover the imaging area. From these results, the 60 × 10 deg LSD is the most preferable for the LD of the HLC under the various assumed usage conditions.

**Supplementary Fig. 5.**
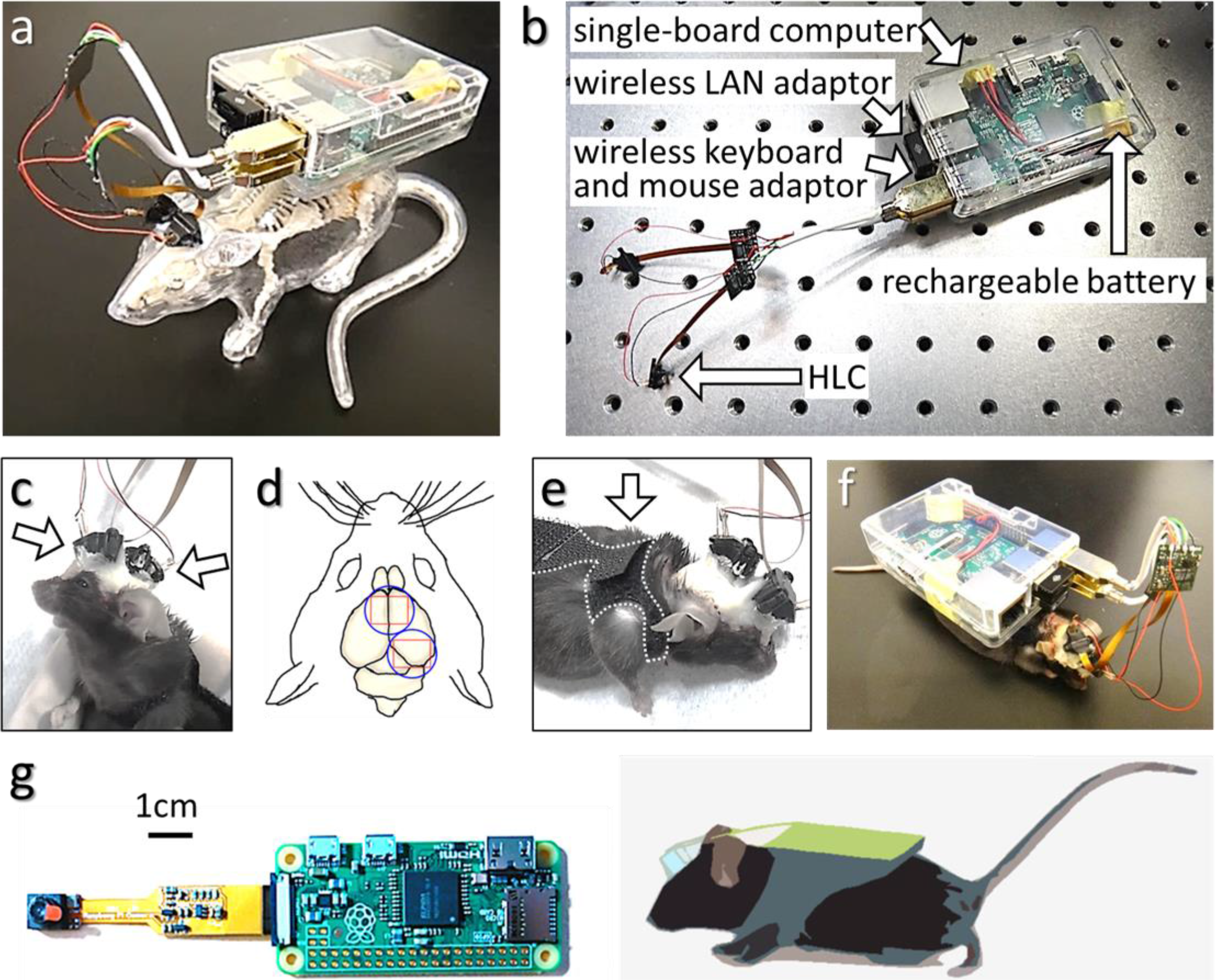
Wearable on-demand wireless imaging system using single-board computers. An on-demand device is preferable for real-time observation of natural behaviors. Therefore, we constructed several types of wearable wireless units by using single-board computers. The CMOS image sensor used in the HLC is more compact and can work on lower power than a CCD (charge-coupled device), and is suitable for wireless control. **(a, b)** Examples of a wearable on-demand wireless imaging system are shown. Dual HLCs were applied to the visual cortex of both hemicephalons on the rodent maquette (a). A Raspberry Pi 2 model B (Raspberry Pi Foundation, UK) was used as a main single-board computer (b). This multi imaging system is composed of the HLCs and adaptors for wireless LAN, keyboard, and mouse. Therefore, it can autonomously perform imaging and recording, and accept control from other PCs as a slave and send real-time video by remote operation. The wireless multi imaging system whose weight is 48 g can run for approximately 6-10 minutes with a rechargeable battery. The operating time can be extended depending on battery capacity. **(c)** An example of multiple uses of the HLC is shown (see for detail Supplementary Fig. 1c, d). **(d)** A schematic image of the applied position of (c) where the frontal or occipital cortical area including the motor or visual cortex is shown. The blue circle and red square indicate the spacer position and the imaging area, respectively. **(e)** The image indicates the mouse wearing a vest to attach the device. In order for the mouse to move freely, it is necessary to carry the unit on its back (see also schematic image in (g)). **(f)** An example of the wireless multiple imaging system (a) applied to a C57BL/6 line mouse. However, the weight and size of this type of system are slightly large and heavy for the tiny body of mouse. For instance, the ICR mice line that has a relatively large body size, a rat, or other large animals will be suitable for this wireless system. **(g)** The system can be scaled down by using Raspberry Pi Zero as shown. Raspberry Pi Zero is smaller and lighter than Pi 2, approximately 1/12 by volume. Therefore, this type of wearable wireless imaging system will be more suitable for small animals such as the C57BL/6 line mouse.

**Supplementary Fig. 6.**
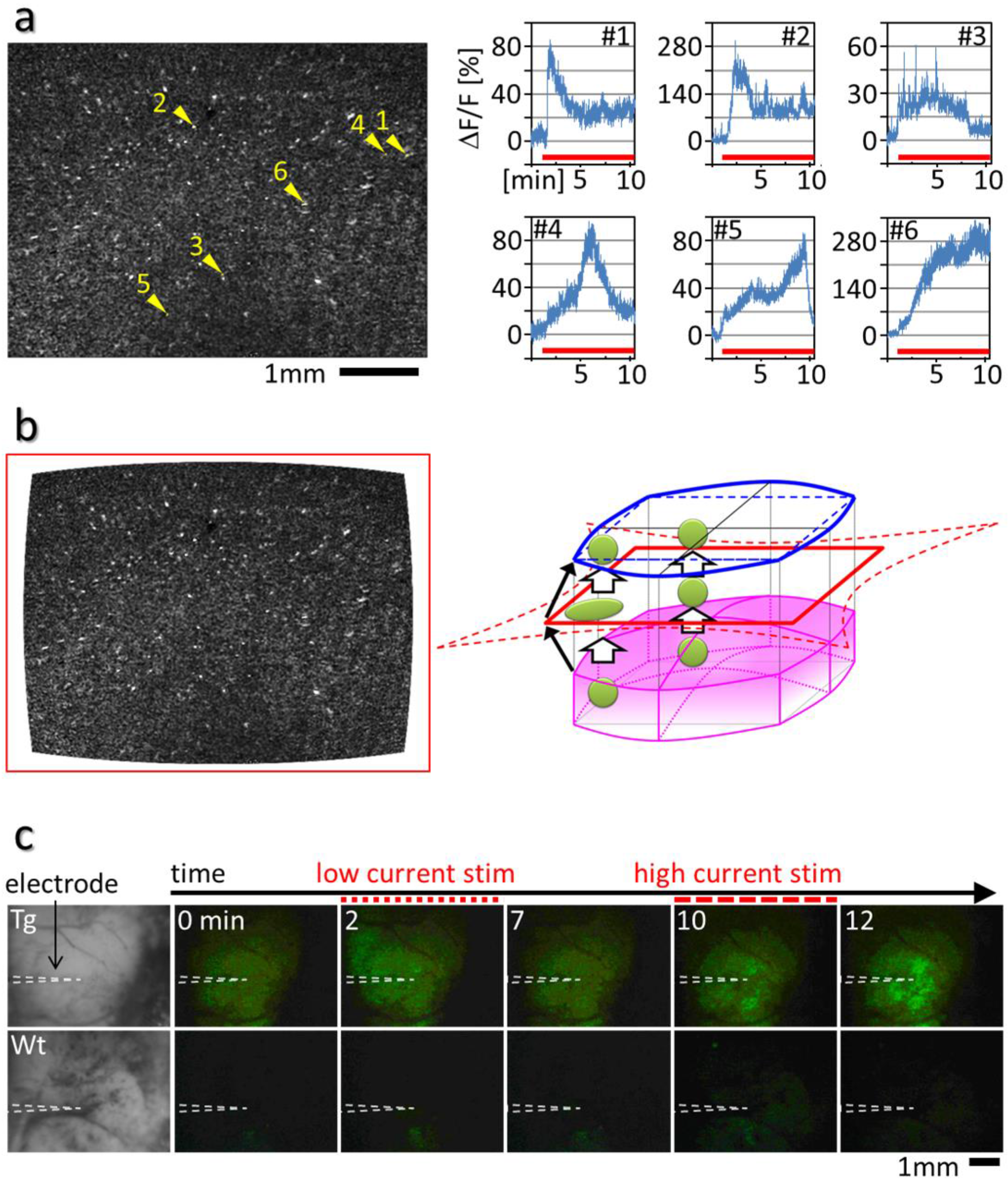
The HLC can visualize intracellular Ca^2+^ dynamics of individual cells in 3D culture and artificially evoked neuronal activities by electrostimulation in the deep cerebral layer. **(a)** The results of individual analysis of the Ca^2+^ imaging by the HLC of the 3D cultured Hela cells that were transfected with the G-CaMP6 gene (Fig. 4a) are shown. The 106 fluorescence spots in the imaging movie (10 frame/sec, 10 min) were selected randomly as a 1 or 4 pixels ROI. The mean of fluorescence intensity (FI) changes at each ROI and the change rate of FI (ΔF/F) was calculated as (F-F_0_)/F, where F_0_ is the mean FI of the first 10 frames in each ROI. The maximum ΔF/F was 693.3 % in these ROIs. The positions of a representative 6 ROIs are shown by the arrowheads with numbers in the image of (a), and their ΔF/F are shown in the graphs and movie (Supplementary Movie 4). The fluorescence intensity was increased after histamine administration (red line in the graph). The graphs are arranged in chronological order of the timing of the ΔF/F peak appearance. The results indicate that the fluorescence functional cell imaging that represent the Ca^2+^ dynamics within each cell can be performed by the HLC at single-cell resolution. For traditional 2D culture conditions without embedding into the extracellular matrix gel, the FI of the Ca^2+^ indicators usually promptly starts to increase just after histamine administration, or within 2 minutes at the latest (supplemental information in the previous study^12^). Under the 3D culture condition, however, the response may be delayed for longer periods depending on individual cells, sometimes as long as more than 10 minutes. Such a delayed response might depend on the depth of each cell because there should be a natural time lag in the diffusion of the histamine into the interior of the gel. **(b)** The left image of (a) was corrected with DTV (television distortion) −3.9 % according to the result of Fig. 2a. Red square line indicates the image silhouette of (a). The right schematic diagram shows the correction process of the image. Magenta block indicates the thick convex image area of the HLC. The magenta gradient schematically represents the difference in fluorescence detection ability according to the result of Fig. 3e. A red line indicates the image silhouette of (a) that is taken by the HLC. The red broken line indicates the positive pincushion distorted actual image of the original object (magenta block). The blue line indicates the negative barrel compensated image of the red line, and also indicates the restored original image silhouette of the object (magenta block). The blue broken line indicates the predicted non-compensated image of the blue line, in other words, the positive barrel distorted image of the blue line. The distortions of the schematic images are enhanced for the ease of visual understanding. Black arrows show the same corner of each image. Green circles indicate the cells. The distortion is smaller near the center of the original object, while the distortion is larger near the edge of the original object. Therefore, the cell shape that was enlarged at the periphery in the image taken by the HLC is corrected to the original size in the restored image (left picture). **(c)** Ca^2+^ imaging was performed by the HLC on the somatosensory cortex of the Thy1-G-CaMP7 transgenic mouse (Fig. 4c, d). We performed Ca^2+^ imaging with the HLC for evoked activity before Ca^2+^ imaging of other physiological neuronal activities. In this experiment, an HLC equipped with 450-460 nm LD, 30 × 5 deg LSD and >520 nm emission filter was used. After craniotomy, a coaxial electrode (TOG207-078, Unique Medical Co., Ltd., Japan) was inserted into the somatosensory cortical layer 5/6 at around 500-600 μm depth using a micromanipulator. Then, a low or high tetanic stimulation (3 or 10 mA biphasic square waves, 100 Hz, duration 150 μs, interval 10 ms, 3 sec.) was applied by using a stimulator (Isolated pulse stimulator 2100, A-M Systems Inc., USA). The time lapse imaging data of the *in vivo* experiment (Fig. 4c, d) on the Thy1-G-CaMP7 mouse (Tg) or wild type mouse (wt) for the negative control are shown. The electrode was inserted into the cortex from the side after the craniotomy. The HLC was attached on the cortex and then the Ca^2+^ imaging was performed. When the stimulation was applied, the FI increased around the electrode in the Tg mouse, and the response was greater after high current than low current stimulation. In contrast, similar evoked signals were not detected in a wt mouse. These results indicate that the HLC can detect evoked neuronal activities in the cortex. Especially, it is thought that these changes of the FI mainly reflect the activities of the layer 5/6 pyramidal neurons and its neurites, because the Thy1 promoter is activated in those neurons predominantly. In the wild-type case, slow and slight increases of the FI were also detected after stimulation. It is thought that this reaction reflects the intrinsic flavoprotein fluorescence in mitochondria^41^. The excitation peak is near 450 nm, therefore, the endogenous flavoprotein imaging depending on neural activity is also accessed by the HLC although the amount of FI change is much smaller, and the response speed is much slower than G-CaMP. Such flavin imaging would allow the functional neuronal imaging in the completely intact brain.

**Supplementary Fig. 7.**
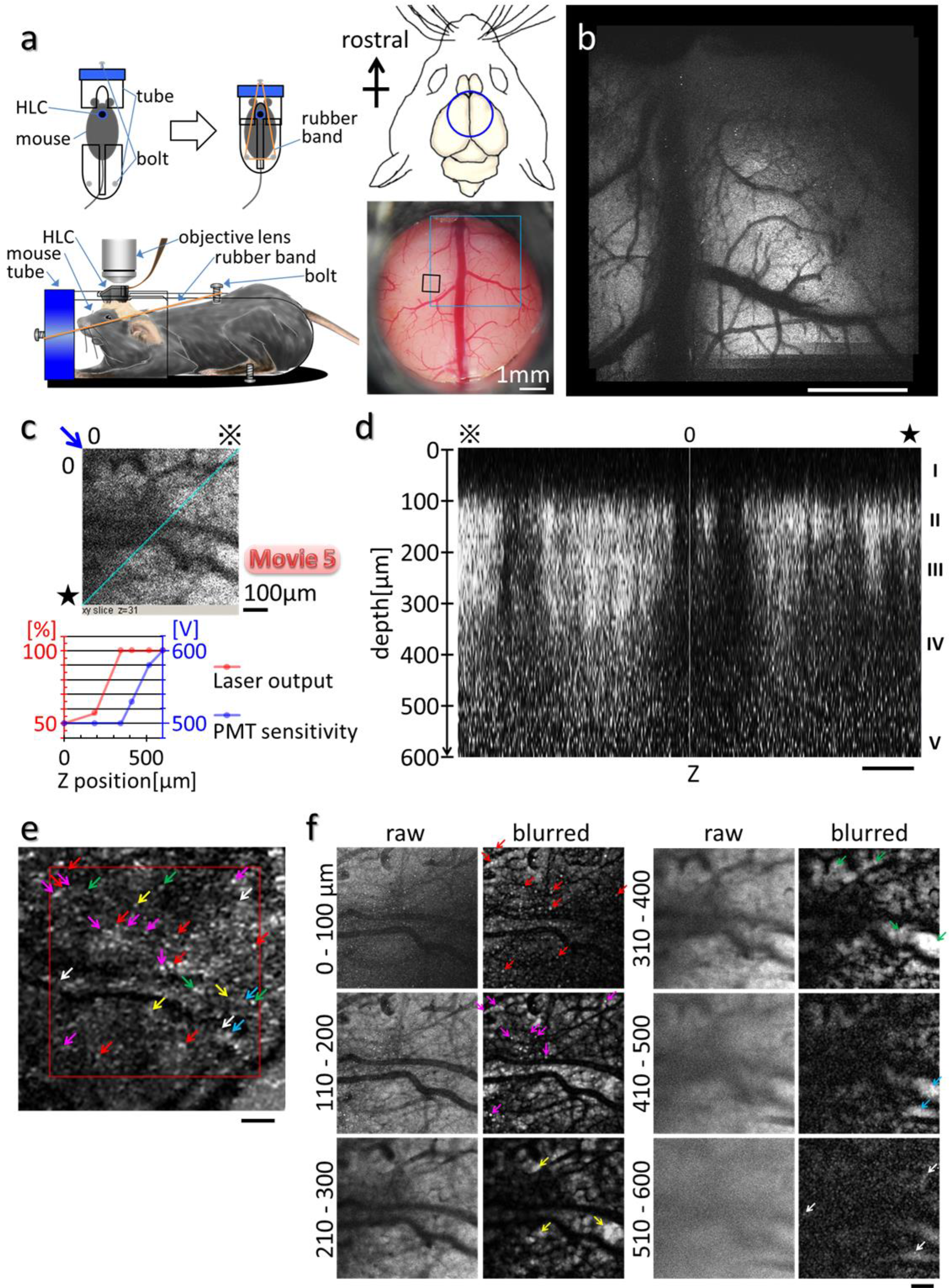
The spacer apparatus with the cranial window allows the HLC and 2-photon microscope to perform chronic Ca^2+^ imaging at the same position in the same mouse. **(a)** Fixation of the mouse head with the HLC on the head. After craniotomy and removal of the dura mater, the CaMK2a-G-CaMP7 mouse (mouse ID: # 3) was installed with the spacer apparatus with a cranial window on the frontal cortical region (see for detail Supplementary Fig. 1a, b). Then, fluorescence imaging was performed by using a 2-photon microscope (FVMPE-RS, Olympus, Co., Japan) at 5-6 days (b) and 33 days (c, d, f) after the surgical operation. The mouse was held in a plastic tube with the head fixed by the attachment of the HLC spacer to the hole of the tube (schematic image of left panel). The schematic image of the right upper panel indicates the location of the cranial window (blue circle). The right lower image was taken by a stereo microscope through the cranial window. Blue or black square indicates the imaging area of (b) or (c, d, f), respectively. **(b)** The fluorescence imaging was performed in the awake condition with a 2-photon microscope using a galvano scanner. The view field was 3.18 × 3.18 mm, 512 x; 512 pixels, 6.21 μm/pixel, and images were captured 10 times at 1.01 sec/stack at the depth of 300 μm. The acquired imaging data were merged. Misalignment caused by mouse movements between stacks was corrected manually, and a “maximum intensity projection”, *i.e.* a creation of the image where each pixel contains the maximum value over all images in the stacks, was performed. The result is shown in (b). Each fluorescent spot should represent excitatory neurons. Bar indicates 1 mm. **(c)** The fluorescence imaging was performed with a 2-photon microscope using the galvano scanner in the anesthetized condition (upper panel). The view field is 636 × 636 μm, 512 × 512 pixels, and the images at 0.917 sec/stack were captured to a depth of 600 μm every 10 μm. The image shows a horizontal section of the cerebral cortex at the 31th stack, that is, a depth of 300 μm. The blue line and blue arrow indicate the position of the vertical section of the cortex and the direction of view in (d). The reference marks of two type asterisks correspond to the positions shown in (d). The lower graph indicates the imaging setting. Depending on the depth, the laser output and PMT (photomultiplier tube) sensitivity of the 2-photon microscope were changed. Time-lapse imaging of the same view area in (c) at a 120 μm depth of the anesthetized mouse cortex was performed with a 2-photon microscope using a resonant scanner. The 10 times speed movie that was captured at 389.72 ms/frame and averaged at every 10 frames is shown in Supplementary Movie 5. Since the resonant scanner allows faster capture than the galvano scanner, the blinking of fluorescence spots is easier to recognize. It is thought that those fluorescence spots blinking represent the activity of excitatory neurons. **(d)** Image shows the vertical cross section of the cortex (c). The length of 0-※ and 0-★ are 900 μm. Roman numerals on the right side of figure indicate the layer of the cortex. Bar indicates 100 μm. **(e, f)** The fluorescence image of the same mouse as (a) which was taken by the HLC under the freely-moving condition at 57 days after surgical operation is shown in (e). Red square shows the imaging area by 2-photon microscopy, which is the same as the black square in (a). Bar indicates 100 μm. Each colored arrow roughly corresponds in position to the light spots in (f). The image of is the result of the processing as the following. The fluorescence movie data taken by the HLC for 5 minutes at 50 ms/frame (20 fps) was deconvoluted and background subtracted, and all frames were merged to one image by a maximum intensity projection. In (f), the fluorescence images of the anesthetized mouse were taken by 2-photon microscopy. Bar indicates 100 μm. The images at 1.00 sec/stack were captured to a depth of 600 μm at every 10 μm intervals (see for the Z axis reconstruction data in (d)). And then, images were superimposed every 100 μm by the maximum intensity projection of each of 10 slices, as shown “raw”. The blurring with a radius of 3.8 pixels at the light spot of that raw imaging data was performed as shown “blurred” by using a Gaussian filter of ImageJ. We performed this blurring procedure to match the resolution (9.62 μm/pixel) by the HLC whose view field in this experiment is 4.62 × 6.16 mm, 480 × 640 pixels to the resolution (1.24 μm/pixel) by 2-photon microscope. The difference in resolution between the HLC and 2-photon microscope is 7.7 times, therefore the radius of 3.8 pixels blurring was performed to obtain similar images. As a result, during the long-term housing, neuronal activity could be visualized using the HLC and also by conventional 2-photon microscopy through the cranial window of the spacer apparatus.

**Supplementary Fig. 8.**
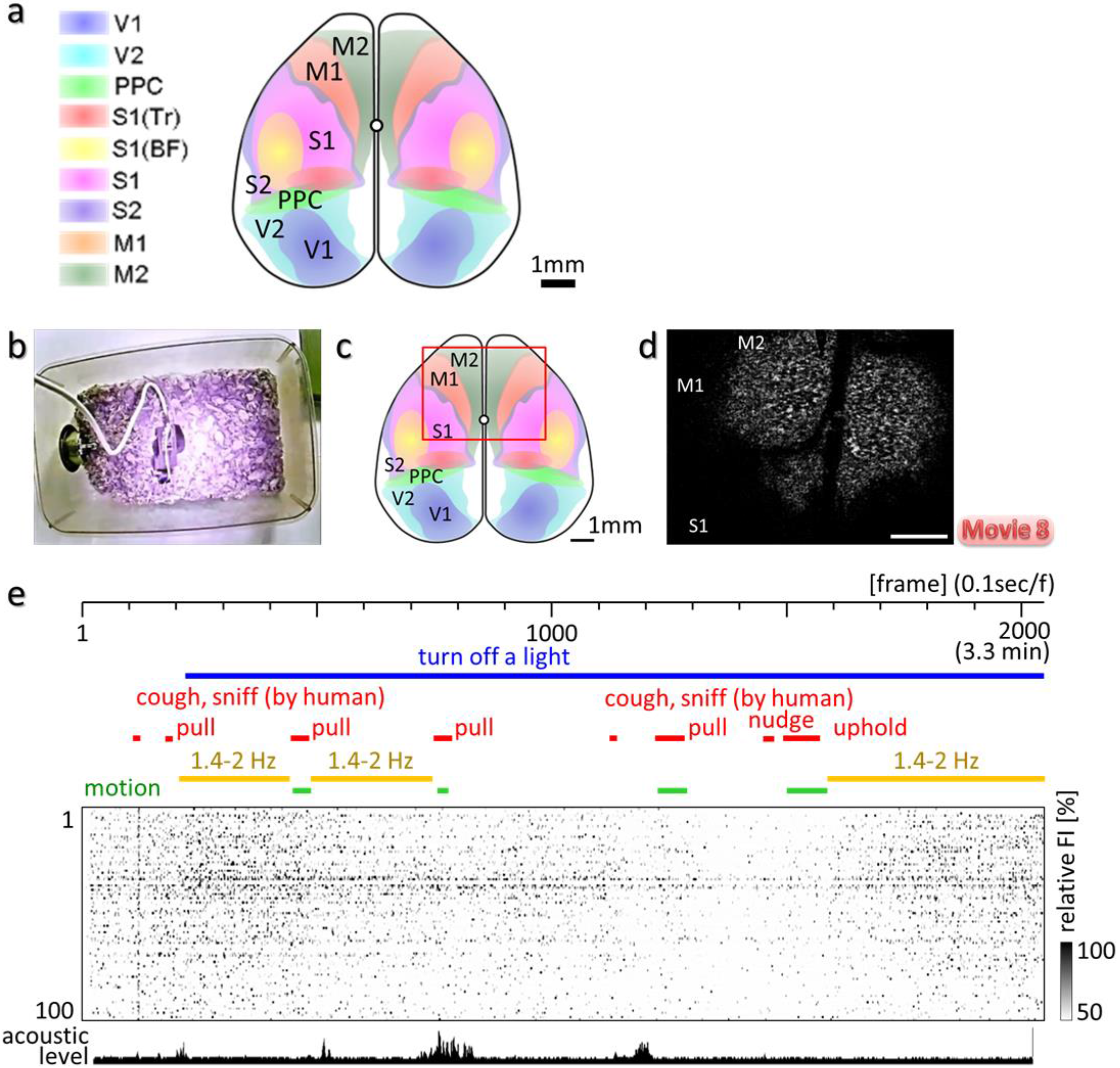
Event related neuronal activity was observed in the frontal cortex in the freely moving condition. Ca^2+^ imaging was performed with the HLC in the freely moving CaMK2a-G-CaMP7 mouse in the square cage. The behavior of the mouse was taken with an IR web camera placed above the cage. **(a)** The schematic image shows the cerebral cortical map which was partially modified from the map^37^ reconstructed from the serial coronal sections of the brain atlas^36^. Each abbreviation means V1/2, primary/secondary visual cortex; PPC (PtA), posterior parietal cortex (parietal association cortex); S1 (Tr/BF)/S2, primary (trunk/olfactory barrel field)/secondary somatosensory cortex; M1/2, primary/secondary motor cortex. **(b)** The image shows the experimental setup. **(c)** The schematic image of the HLC imaging area is shown (red square). **(d)** The subtracted fluorescence image of the frontal cortex (c) in the CaMK2a-G-CaMP7 mouse taken by HLC under the freely moving condition (b) is shown. The raw movie (10 frame/sec) was subtracted by the averaged frame of the first 10 frames of the movie, and then the brightness and contrast was enhanced to observe neuronal activity in association with behavior as shown in (d, Supplementary Movie 8). Bar indicates 1 mm. **(e)** 100 fluorescent spots of 4 pixels were picked up at random as representative ROIs from both hemispheres broadly based on the raw data. A temporal change of the FI in each ROI is shown by the raster plot. Several interventions were applied to the mouse during Ca^2+^ imaging (notation in red). The bottom graph shows an ambient acoustic level. The spacer of the HLC was shielded from light by the additional liquid rubber coat, and the changes in the surrounding lighting had no direct influence on the imaging result. The blue line indicates the period of dark environment, and the mouse behavior could still be observed by the IR web camera. Since the HLC has an IR cut filter (<650 nm), IR lighting also had no direct influence on the imaging result. The green lines indicate mouse motion. When pulling, nudging, and upholding the mouse body were repeatedly conducted during Ca^2+^ imaging of the frontal cortex, neuronal activity changed in relation to these events. When the experimenter coughed near the mouse, transient and overall activity occurred in many ROIs. However, such a global reaction was not caused by a second cough, possibly due to acclimation. Broad slow synchronized wavelike activity propagated sometimes at the resting state during the periods indicated by yellow lines. Such a slow wave response was interrupted by a pull, and does not always occur during the resting period, which suggests that the phenomenon is not just noise. This slow oscillatory activity may be similar to the one observed during the inter trial periods^43^.

**Supplementary Fig. 9.**
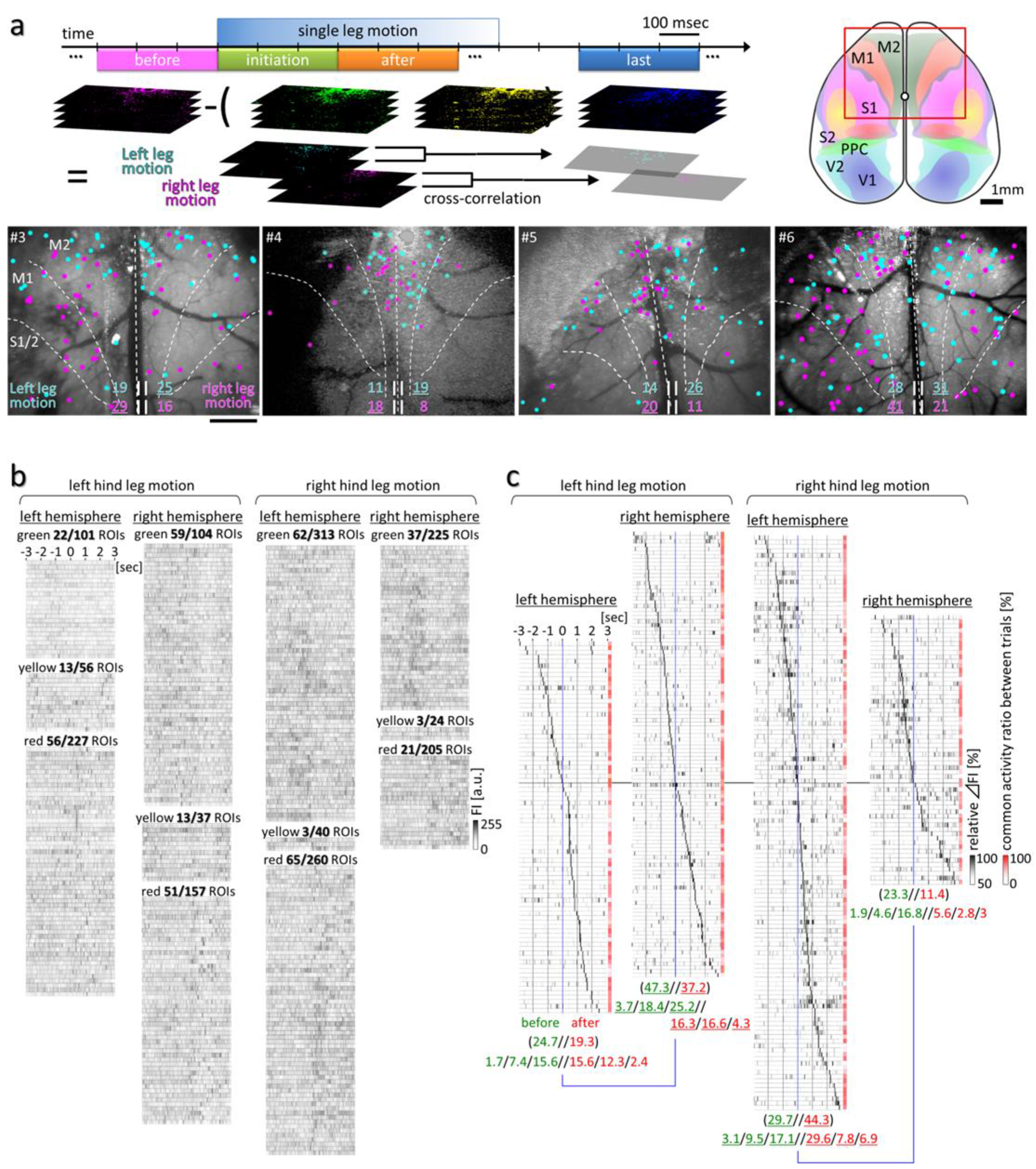
Cross-correlation analysis and the relative ΔFI in the comprehensive quantitative analysis reveal the specific cell assemblies which represent a cascadic premotor activity during voluntary movement. **(a)** The result shows the cross-correlation analysis of premotor activity. In the CaMK2a-G-CaMP7 mouse, 10 or 20 fps (frame per second) Ca^2+^ imaging was performed over the frontal cortex including the whole motor cortex by the restriction motion experiment and the HLC (Fig. 8a). The left schematic image indicates the process of the analysis method. The middle schematic image shows the imaging position by the HLC. The red square indicates the imaging area by the HLC, 5.0 × 6.5 mm. The activity positions of excitatory neurons during the 300 ms before the onset of the left or right hindleg motion were extracted in the process as described below. First, 4 periods for 300 ms during the left or right hindleg’s voluntary movement were categorized as “before”, “initiation”, “after” and “last”. For example, “before” or “initiation” was defined the period just before or after the onset of the motion, respectively. “After” was defined as the period just after “initiation”. “Last” was defined as the period from 200 ms just after the motion stopping. Next, fluorescence imaging data from each of the 4 periods was subtracted by the data of the one frame just before each period to reduce the background. Those subtracted imaging movie data were converted to a one frame image by maximization, and then, the subtraction between each period, max of “before” - max of [“initiation” + “after” + “last”], was performed for each ROI. The result was expected to represent only premotor activity. Finally, the image of two discrete trials accompanying the left or right hindleg’s motion in the same mouse are shown for comparison in the lower images of (a). The results of 4 mice are shown (mouse ID: #1 - #4). Bar indicates 1 mm. Cyan or magenta dots indicate the neurons activated before the left or right hindleg motion. For example in #1 mouse, 19 and 25 neurons of the left and right hemisphere, respectively, were activated before the left leg motion. And 29 and 16 neurons of the left and right hemisphere, respectively, were activated before the right leg motion. These neurons seem to be more localized to M2 than M1, and are rarely present in the other areas such as the somatosensory cortex, and its localization is biased toward the contralateral side over the motion side. The results of (b, c) show a part of the comprehensive quantitative analysis of the premotor activity in Fig. 8. **(b)** The temporal value changes of FI, derived from the raw 20 fps Ca^2+^ imaging data, at ROIs in the subtracted image of Fig. 8e with >5 > ΔF/F are presented by a raster plot. The numbers at the top of each raster plot mean [>5 > ΔF/F ROIs] / [all ROIs]. The ROIs were classified by color codes (green, yellow and red) according to the criteria described in the legend of Fig. 8e. The yellow ROIs are double positive for both green and red characters. **(c)** The data shows the temporal raster plot of relative FI change rates of (b) (relative ΔFI [F]), and an averaged possibility of all kick events at each ROI (red boxes). The relative ΔFI was calculated so that a minimum or maximum value was set to 0 or 100 for each ROI in data of (b). Then, the 50-100 % relative ΔFI in each ROI was presented by the order of the peak point time. The possibility value was calculated based on whether the subtracted FI at each ROI in each kick event was positive (= 1) or negative (= 0). The averaged possibility values at each ROI in all kick events are indicated by red boxes at the right end of the raster plot line. The sum of possible values is indicated by the number under each raster plot. Green or red colored numbers mean the sum before or after the initiation of the motion. The bottom numbers mean the sums within each time period (−3 to −2 / −2 to −1 / −1 to 0 // 0 to 1 / 1 to 2 / 2 to 3 [sec]). Underline means the larger numbers compared to those on the other side of the hemisphere. These results also indicate the tendency that many active ROIs exist in the contralateral hemispheres from the motion side similar to the result in (a). Next, an overall qualitative analysis was performed to eliminate the concern that accidentally large but rare activities might make unduly great contributions by maximization analysis. **(d, e)** Left graphs indicate the averaged total ΔF/F of all ROIs (b) during all left or right kick events. ΔF/F was measured from raw data of each event at the ROIs (b) excluding those of the <50 % possibility and the somatosensory area. Then, the ΔF/F of 11 or 10 trials were all averaged for each hemisphere. The middle or right graphs indicate the averaged total ΔF/F of the specific cell assembly whose peak of ΔF/F appeared before or after the initiation of the leg movement (abbreviated as “before group”, “after group”). Bold red and blue lines indicate the mean of FI of each 5 frames. The bottom numbers indicate the numbers of the ROIs in the left or right hemisphere within each time period (−3 to −2 / −2 to −1 / −1 to 0 // 0 to 1 / 1 to 2 / 2 to 3 [sec]). The underlines mean that the number is larger than on the other side. As a result, the lateralized activity was observed, especially at a period from −2 to +2 sec, just before and after the initiation of the motion in left graphs. These three differential activities are also basically observed in the middle graphs representing the result of the “before group” cell assembly. Surprisingly, however, in the “after group” cell assemblies (right graphs), there was almost no difference in the left-right activities at periods from 0 to +2 sec. In contrast, the “before group” cell assemblies (middle graphs) indicate the different left-right activities before and after the onset of the motion. In conclusion, our data suggest that motor planning for the left or right hindleg specific movements has been already completed by the lateralized activation of the “before group” cell assembly before the onset of the movement which lasts even after the initiation of the movements. **(f, g)** The “before group” and “after group” cell assemblies of (d, e) were plotted on the schematic diagram of the motor cortical area whose regional compartments are the same as the image of Fig. 8d. The bottom numbers mean the numbers of the ROIs in the left or right hemisphere. The underlines mean that the number is larger than on the other side. As a result, the tendency of the lateralized distribution also appears in the overall qualitative analysis similarly to visual judgment, cross-correlation analysis, and comprehensive quantitative analysis. In summary, lateralized premotor activities seem to start from M2 on each opposite side of the hemisphere, and the activities propagate to M1 and the somatosensory region. The trends of our results are largely consistent among all the different analyses that were conducted in this paper. The result that the volitional movement originates from M2 is consistent with previous reports and supports them^18,44–46^. The pre- and post-motor activities in M1 have been occasionally pointed out to be non-lateralized^17,47,50^, and even ipsilateral^49,50^. In fact, the “after group” cell assemblies of (d, e) show bilateral non-lateralized activity during the period from 0 to +2 sec. However, the “before group” cell assemblies of (d, e) represent differential activity, and the 2D distribution study in Fig. 8f showed that the pre- and post-motor activities in M1 are also lateralized similarly as well as in M2. Therefore, we conclude that M1 activity is lateralized. The lateralized premotor activity at just before the initiation of the motion (−0.5 to 0 sec) probably represents the start of a direct motor command in the neurons projecting to motor neurons controlling the muscle, and the period includes a point of no return in vetoing the motion (around −0.2 sec)^51^. The direct activity in M1 occurs 50 to 80 ms before muscle movement^52^.

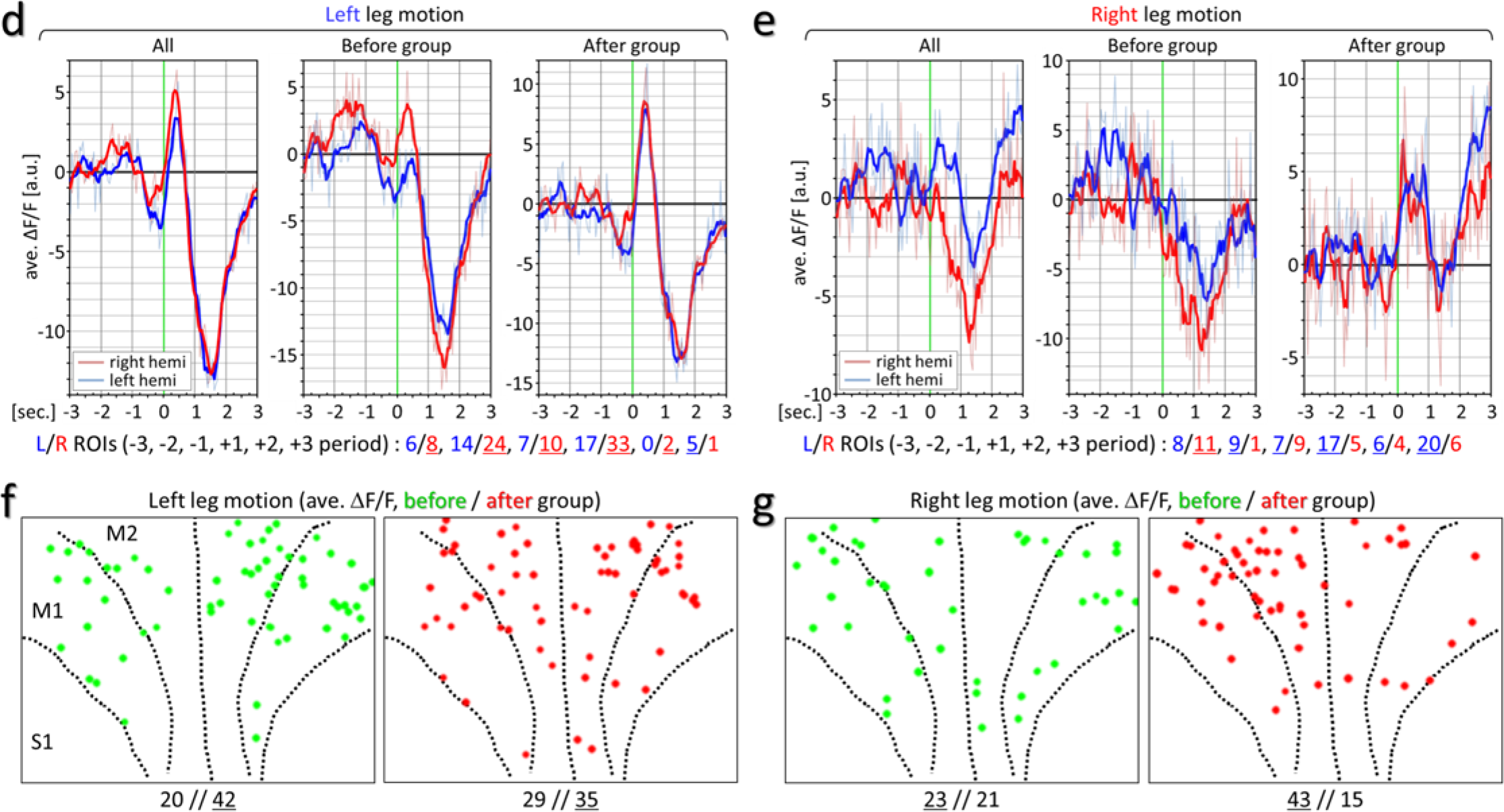

**Supplementary Movie 1** Examples of imaging the freely moving mouse wearing the HLC and spacer apparatus (Fig. 1c) under various conditions.

**Supplementary Movie 2** Ca^2+^ imaging by the HLC over the occipital cortex of an awake CaMK2a-G-CaMP7 mouse (Fig. 1d).

In the movie, (a) shows a movie where the background of the raw imaging movie (10 frame/sec) was subtracted by using ImageJ with a sliding paraboloid (curvature radius is 200 pixels), and was deconvoluted. (b) is one frame of the movie (a). (c) is the magnified movie of the inset shown in (b). (d) is the same as (b). (e) was made by subtracting the FI of each pixel with the average of the FI of the same pixel during the first 10 times.

**Supplementary Movie 3** The procedure for counting the number of blinking spots in the view field of Fig. 1d.

In the movie, (a) shows the maximized movie that 8 movies of 1 min were merged by taking the maximum FI for each pixel. (b), (c) show the same image of Supplementary Fig. 3a, b, respectively.

**Supplementary Movie 4** x10 Ca^2+^ imaging by the HLC on the 3D cultured G-CaMP6-Hela with histamine administration (Supplementary Fig. 6a, raw fluorescence image is shown in Fig. 4a).

**Supplementary Movie 5** x20 Ca^2+^ imaging with a 2-photon microscope over the motor cortex of the anesthetized CaMK2a-G-CaMP7 mouse (Supplementary Fig. 7c).

**Supplementary Movie 6** x100 Ca^2+^ imaging by the HLC on the somatosensory cortical area of the anesthetized Thy1-G-CaMP7 mouse (Fig. 4d, raw fluorescence time-lapse image is shown in Supplementary Fig. 6c).

**Supplementary Movie 7** Magnified Ca^2+^ imaging by using the HLC with short spacer in the freely moving mouse.

Ca^2+^ imaging of the visual cortex of the CaMK2a-G-CaMP7 mouse was performed by the HLC with the short spacer under the freely moving condition. The 20-fps movie was magnified by the optical zoom (x 2 in comparison with Supplementary Fig 3) and the digital zoom (x 4) is shown (Fig. 5).

**Supplementary Movie 8** The Ca^2+^ imaging by the HLC of the frontal cortex in the freely moving CaMK2a-G-CaMP7 mouse (Supplementary Fig. 8b, d).

The left and right movies show a behavior movie of the mouse and the subtracted and averaged Ca^2+^ imaging movie, respectively.

**Supplementary Movie 9** x1/10 Ca^2+^ imaging by the HLC on the motor cortical area of the awake CaMK2a-G-CaMP7 mouse (Fig. 8b).

Left/ middle/ right movies show a behavior movie of the mouse, the raw Ca^2+^ imaging movie, and the marked raw Ca^2+^ imaging movie, respectively. In the right movie, the magenta or green dots mean the neuronal activities which start during the before (−4 to 0 sec) or after (0 to +4 sec) the initiation of the motion. Several arrows indicate the dots and its corresponding positions.

**Supplementary Movie 10** Ca^2+^ imaging by the single and dual HLC under the freely moving condition.

Real-time Ca^2+^ imaging of the awake CaMK2a-G-CaMP7 mouse by the single HLC of the frontal cortical area (Supplementary Fig. 2d) or by the dual HLC of the frontal and occipital cortical area (Supplementary Fig. 1d) were performed. The movie shows 30-fps captures of the video images displayed on the PC monitor. Judging from the orientation position of the blood vessel, the vibration induced by mouse’s movements is not recognized and the view field does not change. Also, even if the two HLCs are used, the movement of the mouse is not impaired.

Author Contributions
H.O. and T.K. conceived and designed the research, and wrote the paper. T.K. constructed the imaging system and the experimental apparatus, and performed all experiments. T.I. performed NMF. M.S., M.O., J.N. and Y.H. made the G-CaMP7 transgenic mouse lines.

